# TRAFICA: An Open Chromatin Language Model to Improve Transcription Factor Binding Affinity Prediction

**DOI:** 10.1101/2023.11.02.565416

**Authors:** Yu Xu, Chonghao Wang, Ke Xu, Yi Ding, Aiping Lyu, Lu Zhang

## Abstract

In silico transcription factor and DNA (TF-DNA) binding affinity prediction plays a vital role in examining TF binding preferences and understanding gene regulation. The existing tools employ TF-DNA binding profiles from *in vitro* high-throughput technologies to predict TF-DNA binding affinity. However, TFs tend to bind to sequences in open chromatin regions *in vivo*, such TF binding preference is seldomly considered by these existing tools. In this study, we developed TRAFICA, an open chromatin language model to predict TF-DNA binding affinity by integrating the characteristics of sequences from open chromatin regions in ATAC-seq experiments and *in vitro* TF-DNA binding profiles from high-throughput technologies. We applied self-supervised learning to pre-train TRAFICA on over 13 million nucleotide sequences from the peaks in ATAC-seq experiments to learn the TF binding preference *in vivo*. TRAFICA was further fine-tuned using the TF-DNA binding profiles from PBM and HT-SELEX technologies to learn the association between TFs and their target DNA sequences. We observed that TRAFICA significantly outperformed both machine learning-based and deep learning-based tools in predicting *in vitro* and *in vivo* TF-DNA binding affinity. These findings indicate that considering the characteristics of sequences from open chromatin regions could significantly improve TF-DNA binding affinity prediction, particularly when limited TF-DNA binding profiles from high-throughput technologies are available for specific TFs.

## 1 Introduction

The binding of transcription factors (TFs) to DNA is a crucial step in gene regulation, where TFs recognize and interact with specific promoter sequences in genomes to regulate the expression of nearby downstream genes. The *in vitro* TF-DNA binding affinity is primarily attributed to intrinsic TF binding preferences, which can be captured through *in vitro* high-throughput experiments in the absence of cellular environments [1]. The *in vivo* TF-DNA binding affinity is more intricate, because it not only relates to intrinsic TF binding preferences but is also influenced by other factors, such as the positions of nucleosomes and the co-binding mechanism [2, 3]. Nucleosome positions can restrict the accessibility of TF binding sites, thus TFs could only bind to DNA sequences in open chromatin regions; the co-binding mechanism refers to the cooperative binding of multiple TFs and co-factors, such as the transcriptional corepressor TUP1-SSN6 complex in yeast [4].

Several technologies have been developed to measure TF binding preferences, both *in vivo* and *in vitro*. Chromatin ImmunoPrecipitation followed by sequencing (ChIP-seq) [5] is a popular *in vivo* sequencing technology to collect the binding sequences of a specific TF across the whole genome. ChIP-seq has a significant limitation, as the antibodies for chromatin immunoprecipitation may not always be available for many TFs [2]. Protein Binding Microarray (PBM) [6] is an *in vitro* high-throughput technology that measures the binding affinities of a specific protein to a set of artificial DNA sequences according to fluorescence intensities. High-Throughput Systematic Evolution of Ligands by EXponential enrichment (HT-SELEX) [7] is a well-established *in vitro* technology, which involves multi-cycle selection to screen sequences binding to a target protein. This technology could yield abundant sequences with high affinity to a TF of interest. Moreover, Assay for Transposase-Accessible Chromatin using sequencing (ATAC-seq) [8] is designed to capture chromatin accessibility (**Figure 1A**), which is a key factor that could influence *in vivo* TF binding. The intrinsic preferred sequences (e.g., promoter and enhancer sequences) of TFs are also typically found in open chromatin regions [9]. The rapid development of these technologies has contributed to large-scale publicly available datasets that have yet to be fully utilized [10, 11, 12].

**Figure 1.**
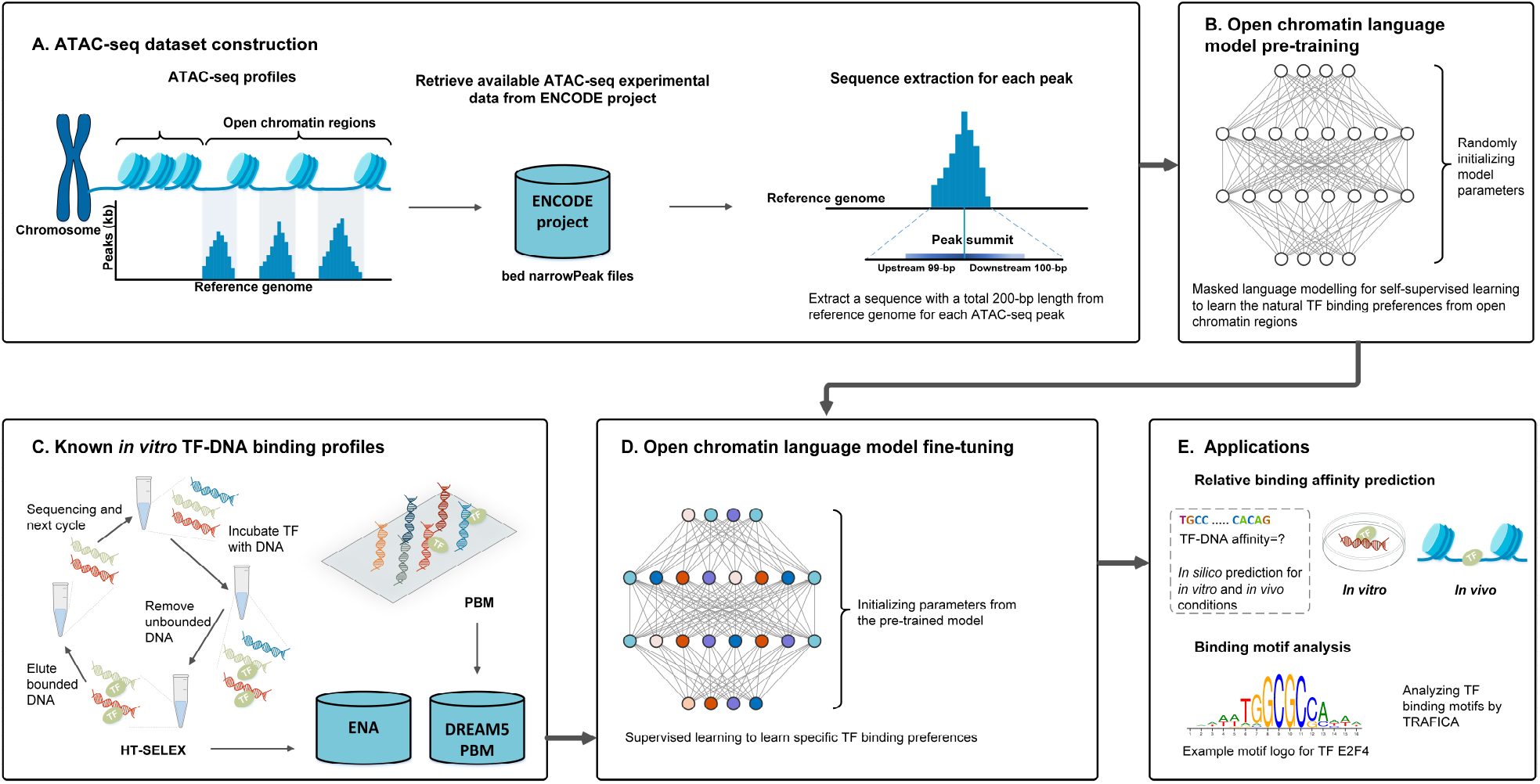
Overview of TRAFICA. **(A)** Extract sequences from open chromatin regions in the human reference genome for TRAFICA pre-training. The open chromatin regions were collected from the narrowPeak files of ATAC-seq experiments in the ENCODE project [10, 11]. **(B)** Pre-training stage of TRAFICA using sequences from open chromatin regions. **(C)** Collection of *in vitro* TF-DNA binding profiles from high-throughput technologies. We collected PBM and HT-SELEX experimental data from the DREAM5 PBM study [19] and European Nucleotide Archive (ENA) [12], respectively. **(D)** TRAFICA is fine-tuned on *in vitro* TF-DNA binding profiles. **(E)** Applying TRAFICA to predict *in vitro* and *in vivo* TF-DNA binding affinity and analyze TF binding motifs.

In recent years, there has been a growing interest in developing machine learning-based models [13, 14, 15, 16, 17] to predict TF-DNA binding affinity using TF-DNA binding profiles generated by PBM and HT-SELEX experiments. DNAshapeR [13] employs a multiple linear regression model with *L*2 penalty and utilizes 13 DNA shape features (including inter-base pair types, intra-base pair types, and minor groove width) from DNA sequences to predict TF-DNA binding affinity. DNAffinity [14] represents a sequence as a feature vector composed of 4 classes of DNA properties (base pair parameters, stiffness, sequence pattern, and electrostatics) and employs a random forest regression model for TF-DNA binding affinity prediction. However, machine learning-based tools typically require feature engineering on DNA sequences to obtain fixed-length features to describe the characteristics of nucleotide sequences. In contrast, deep learning-based models could directly learn characteristics from sequences with different lengths instead of relying on feature engineering. DeepBind [15] applies convolutional neural networks (CNNs) to capture binding motifs by CNN kernels and predict TF binding preferences. DLBSS [16] integrates 4 DNA shape features as inputs of CNNs for TF-DNA binding affinity inference. CRPTS [17] combines a long short-term memory network with CNNs as a hybrid model to improve predictive performance. Although the existing tools have demonstrated advancements in predicting *in vitro* TF-DNA binding affinity, they have yet to consider the characteristics of DNA sequences from open chromatin regions.

In this study, we developed TRAFICA (**TR**anscription factor binding **AF**finity prediction **I**ntegrating **C**hromatin **A**ccessibility), an open chromatin language model that integrates the characteristics of DNA sequences from open chromatin regions and *in vitro* TF-DNA binding profiles from high-throughput technologies to predict TF-DNA binding affinity. TRAFICA is built on a transformer-encoder architecture [18], which includes a pre-training stage to capture contextual characteristics of DNA sequences from open chromatin regions based on over 13 million sequences from ATAC-seq data (**Figure 1A, B** and **Figure 2A**). We further collected *in vitro* TF-DNA binding profiles generated from PBM and HT-SELEX experiments for model fine-tuning (**Figure 1C, D** and **Figure 2B**) and evaluation (**Figure 2C**). We compared the performance of TRAFICA with both machine learning-based (DNAshapeR [13] and DNAffinity [14]) and deep learning-based (DeepBind [15], DLBSS [16], and CRPT/CRPTS [17]) tools for TF-DNA binding affinity prediction. We observed that TRAFICA showed superior performance compared with these tools, achieving the highest average Pearson correlation coefficient (PCC=0.649) on the PBM datasets (**Figure 3A**), and the highest average coefficient of determination (*R*^2^=0.972) on the HT-SELEX datasets (**Figure 4A**). We also examined how the size of TF-DNA binding profiles for model fine-tuning influenced the prediction of *in vitro* TF-DNA binding affinity. The results showed that the performance of TRAFICA was robust even with the small numbers of sequences used for model fine-tuning (**Figure 4B**). Moreover, we assessed the performance of TRAFICA in predicting *in vivo* TF-DNA binding affinity by evaluating it on ChIP-seq datasets. The *in vivo* prediction results indicated that TRAFICA outperformed the other tools, especially with limited fine-tuning data (**Figure 5B**).

**Figure 2.**
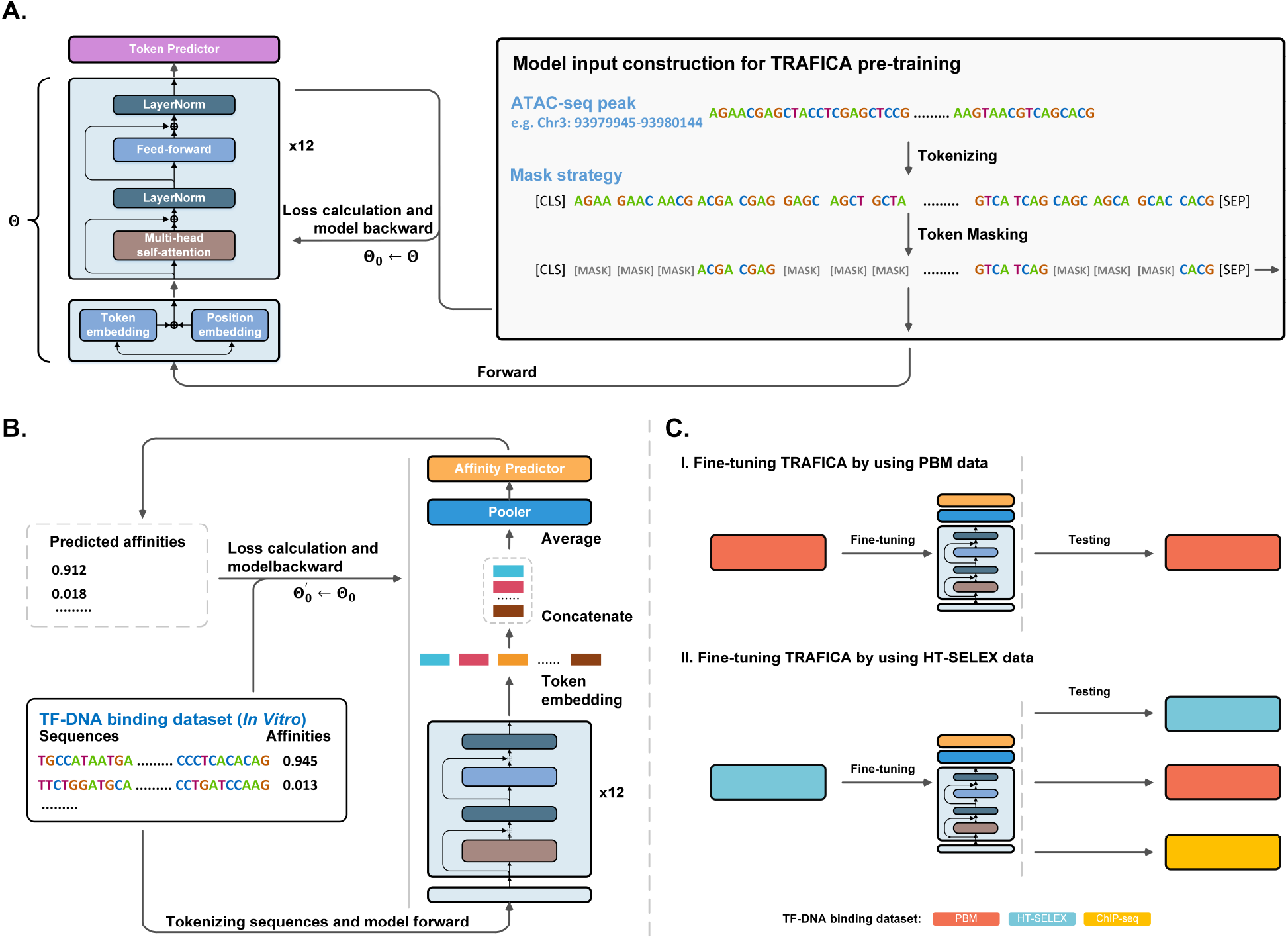
Architecture of TRAFICA. **(A)** Architecture of pre-training module in TRAFICA. The inputs are constructed by tokenizing sequences from open chromatin regions and masking contiguous tokens within them. A token predictor is designed to perform self-supervised learning by predicting those masked tokens according to upstream and downstream unmasked tokens. Θ and Θ_0_ represent the model parameters of the initial and pre-trained models, respectively. **(B)** Architecture of fine-tuning module in TRAFICA. The pre-trained model, coupled with a pooler mod-ule and an affinity predictor, is fine-tuned on TF-DNA binding datasets. Θ_0_ represents the parameters of the fine-tuned model. **(C)** Performance evaluation of TRAFICA. (I) TRAFICA was fine-tuned and evaluated using PBM datasets. (II) TRAFICA was fine-tuned by an HT-SELEX dataset, and its performance was evaluated on the test sets from HT-SELEX, PBM, and ChIP-seq.

**Figure 3.**
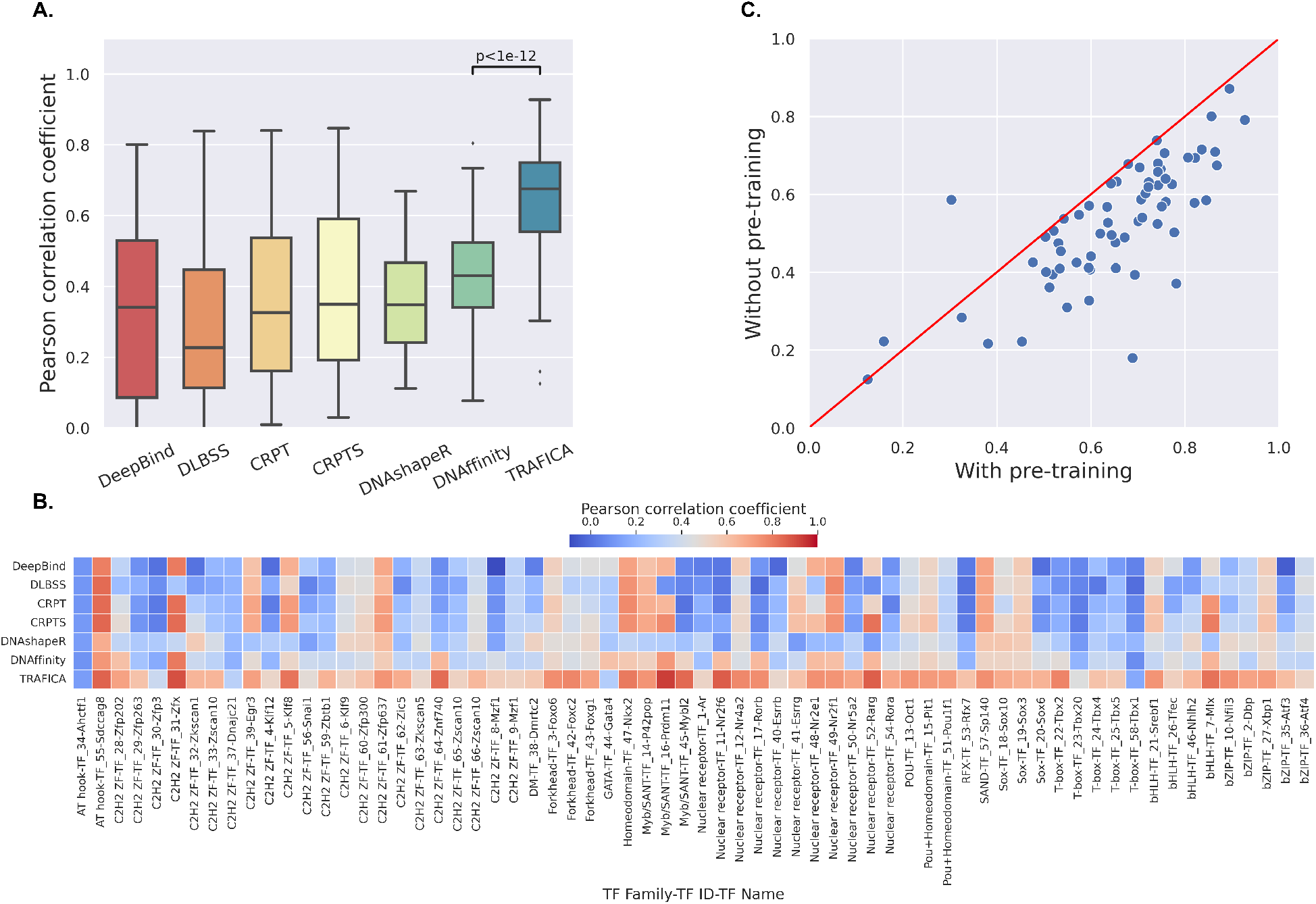
The performance of TRAFICA in predicting *in vitro* TF-DNA binding affinity using DREAM5 PBM datasets. **(A)** Predictive performance of TRAFICA and the other state-of-the-art tools on the PBM datasets. The *p*-values were calculated by Wilcoxon rank-sum test. The black lines inside each box represent the median PCC of 66 TFs. **(B)** Predictive performance of each TF. The *x*-axis labels show the information of 66 TFs, including TF family name, TF ID (defined by the DREAM5 PBM study [19]), and TF symbol (separated by “-”). **(C)** The comparison of PCC values generated by TRAFICA with and without the pre-training stage.

**Figure 4.**
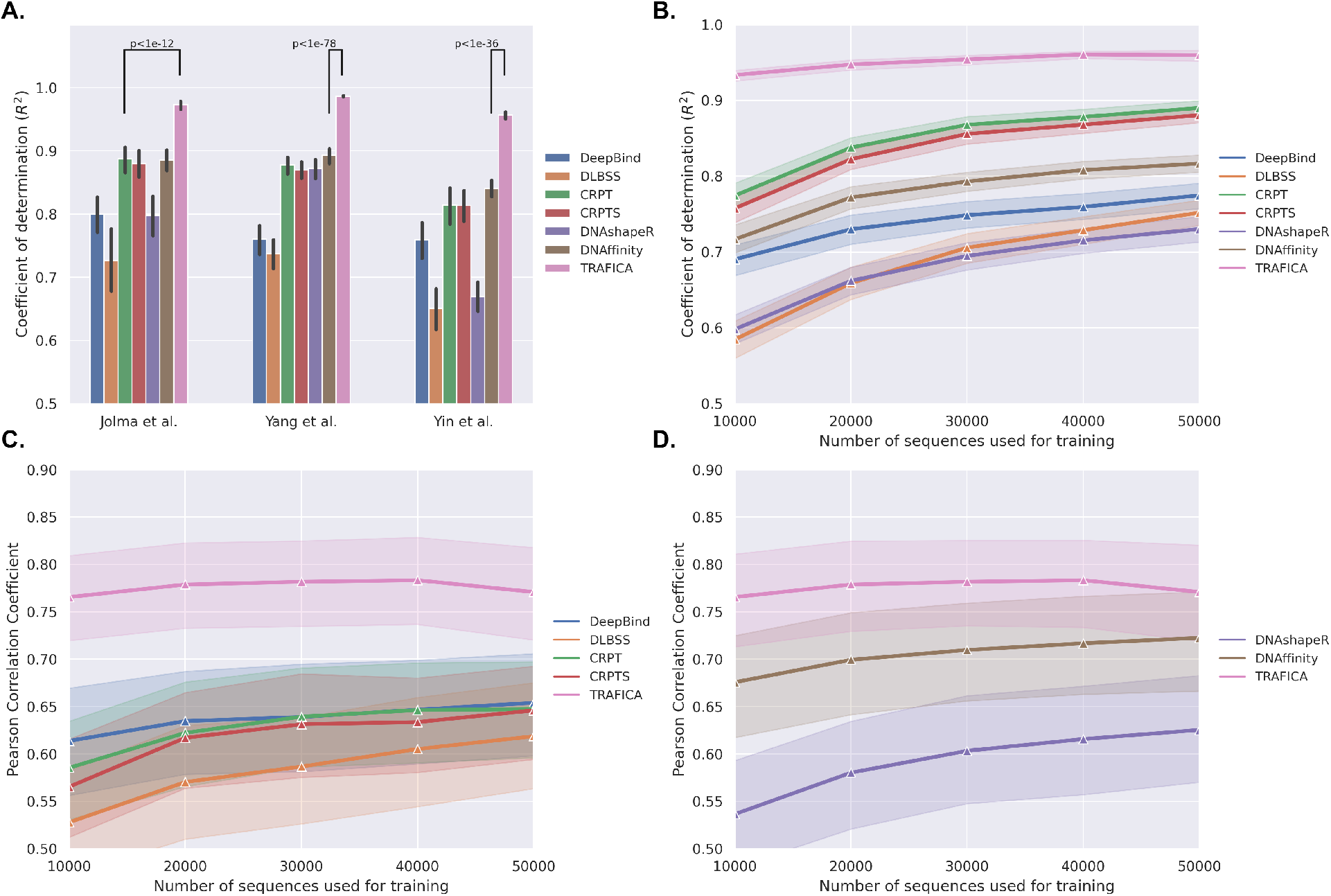
The performance of TRAFICA in predicting *in vitro* TF-DNA binding affinity using HT-SELEX datasets. **(A)** Predictive performance of TRAFICA on the 440 datasets from three HT-SELEX studies. The *p*-values were calculated using Wilcoxon rank-sum test. Error bars represent 95% confidence intervals. **(B)** Predictive performance of TRAFICA (the other tools) using different numbers of sequences in fine-tuning (training) sets on 440 datasets. The shadow areas represent 95% confidence intervals. **(C)** Predictive performance of deep learning-based tools on cross-dataset experiments. We generated different numbers of sequences in fine-tuning (training) sets with only TFs that exist in multiple datasets. **(D)** Predictive performance of TRAFICA and machine learning-based tools (DNAshapeR and DNAffinity) on cross-dataset experiments using different sizes of fine-tuning (training) sets, where the lengths of target sequences are the same.

**Figure 5.**
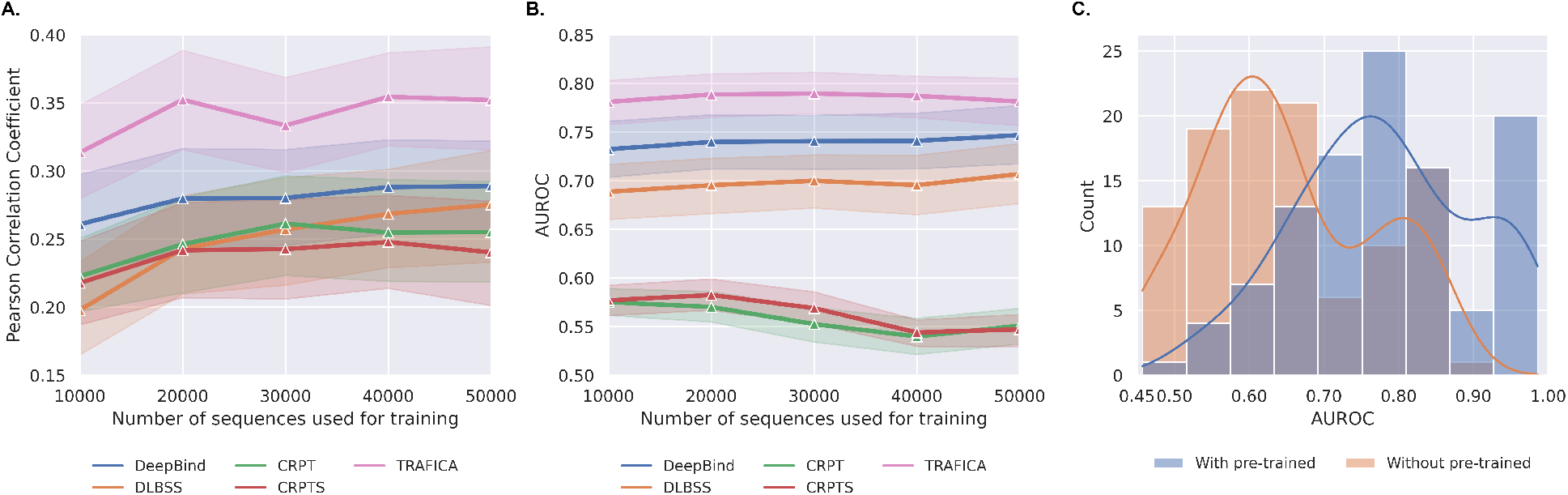
The TF-DNA binding affinity prediction on datasets from different technologies. (**A**) HT-SELEX datasets for model fine-tuning (training) and DREAM5 PBM datasets for model testing. (**B**) HT-SELEX datasets for model fine-tuning (training) and ChIP-seq datasets for model testing. (**C**) Comparing TRAFICA with and without the pre-train stage. 10,000 sequences were used for model fine-tuning.

## 2 Results

### 2.1 Overview of TRAFICA

TRAFICA is an attention-based open chromatin language model that predicts relative binding affinity of particular TFs to DNA sequences by integrating the characteristics of sequences from open chromatin regions measured by ATAC-seq and *in vitro* TF-DNA binding profiles measured by PBM and HT-SELEX technologies. We initially constructed a comprehensive ATAC-seq dataset, which included 13,013,152 genomic sequences from open chromatin regions in the human genome (**Figure 1A**). The open chromatin regions were defined based on the peaks in 422 ATAC-seq experiments (**Supplementary Figure S1** and **Supplementary Table S1**) from the Encyclopedia of DNA Elements (ENCODE) project [10, 11]. These genomic sequences were collected for TRAFICA pre-training to learn the characteristics of sequences from open chromatin regions with natural TF binding preferences (**Figure 1B**). As shown in **Figure 1C and D**, we collected *in vitro* TF-DNA binding profiles measured by PBM and HT-SELEX technologies to fine-tune TRAFICA (PBM: 132 datasets for 66 TFs, **Supplementary Table S2**; HT-SELEX: 440 datasets for 336 TFs, **Supplementary Table S3** and **Supplementary Note 1**). The fine-tuned models were used to predict the *in vitro* and *in vivo* TF-DNA binding affinity, followed by analyzing TF binding motifs (**Figure 1E**).

The architecture of TRAFICA is built on the vanilla transformer-encoder structure [18, 20], which utilizes the selfattention mechanism to capture contextual relationships in sequential data. Each genomic sequence from open chromatin regions is tokenized into ordered *k*-mer tokens as inputs of TRAFICA (**Method**). The framework of TRAFICA consists of a token embedding layer, a position embedding layer, and twelve transformer-encoder blocks (left panel of **Figure 2A**). The two embedding layers are fully connected neural networks designed to encode the semantic and positional information of *k*-mer tokens. Each transformer-encoder block consists of four components, including a multi-head self-attention module, a feed-forward module, layer normalization, and residual connections. Within the multi-head self-attention module, the self-attention mechanism enables the model to capture the complex relationships of *k*-mer tokens in an input sequence by computing attention scores of each token pair, and the multi-head mechanism allows the model to attend to the information of the input sequence more comprehensively. The feed-forward module is a stack of two fully connected layers with a non-linear activation function Gaussian Error Linear Units (GELU) [21], enabling the model to learn intricate dependencies between tokens. Layer normalization is applied to normalize the hidden representation, and residual connections preserve information from earlier layers. The model configuration of TRAFICA is included in **Supplementary Table S4**. Moreover, a token predictor (**Supplementary Figure S2A**) is incorporated to perform masked language modeling in the pre-training stage (**Figure 2A**). This token predictor comprises two fully connected layers, a GELU activation function, and layer normalization. It is discarded after pretraining. Additionally, a pooler module and an affinity predictor (**Supplementary Figure S2B and C**) are employed to predict TF-DNA binding affinity in the fine-tuning stage (**Figure 2B**). This pooler module is implemented as a fully connected layer with a Tanh activation function, and the affinity predictor consists of two fully connected layers. More details about the network structure of TRAFICA can be found in **Supplementary Note 2**.

### 2.2 *In vitro* TF-DNA binding affinity prediction on the DREAM5 PBM datasets

We utilized the DREAM5 PBM datasets to evaluate the performance of TRAFICA in predicting *in vitro* TF-DNA binding affinity (**Figure 2C(I)**). Two microarrays were alternately used as the fine-tuning and test sets for model evaluation: 1. TRAFICA was fine-tuned using the first 33 TFs from the first microarray, and tested on the same TFs from the second microarray; 2. TRAFICA was fine-tuned using the last 33 TFs from the second microarray and tested on the same TFs from the first microarray. We compared the performance of TRAFICA, DeepBind, DLBSS, CRPT, CRPTS, DNAshapeR, and DNAffinity using PCC. This is because the values of relative affinities between the two microarrays were incomparable (**Supplementary Note 3, Supplementary Figure S3**). TRAFICA achieved the best performance (average PCC=0.649), which significantly outperformed the second-best tool DNAffinity (average PCC=0.433, *p*-value<1e-12) (**Figure 3A**). The performance of TRAFICA was more stable than the other deep learning-based tools (standard deviation: 0.159 for TRAFICA; 0.245 for DeepBind; 0.212 for DLBSS; 0.236 for CRPT; 0.235 for CRPTS). We also observed that the predictive performance of all tools varies substantially across different TFs, even within the same TF families (**Figure 3B**). This observation was consistent with the findings in the original DREAM5 study on PBM datasets [19]. Moreover, we performed an ablation study to investigate if including the pre-training stage could improve the performance of TRAFICA. The results indicated a substantial improvement by involving the pre-training stage, with the average PCC rising from 0.526 to 0.649 (*p*-value<1e-5). This highlighted the necessity of integrating natural TF binding preferences as prior knowledge in predicting TF-DNA binding affinity (**Figure 3C**).

### 2.3 *In vitro* TF-DNA binding affinity prediction on the HT-SELEX datasets

For each of the 440 HT-SELEX datasets, the nucleotide sequences were split into a fine-tuning set (80%) and a test set (20%), respectively (**Figure 2C(II)**). The fine-tuning set for each HT-SELEX dataset was employed to train the other tools, respectively. We adopted *R*^2^ to evaluate the performance of TRAFICA because the sequences for model fine-tuning and testing were from the same HT-SELEX dataset (**Supplementary Note 3**). We found TRAFICA substantially outperformed the existing tools on the datasets from all three previous studies (**Figure 4A**). It achieved impressive average *R*^2^ values of 0.973 (Jolma et al.), 0.987 (Yang et al.), and 0.957 (Yin et al.) and substantially better than the second-best tools (Jolma et al.: CRPT, *p*-value<1e-12; Yang et al.: DNAffinity, *p*-value<1e-78; Yin et al.: DNAffinity, *p*-value<1e-36). We also examined if the number of sequences in the fine-tuning set for TRAFICA (training set for the other tools) would influence the performance of these prediction tools by sampling 10,000 to 50,000 sequences (**Methods**). We found the performance of TRAFICA was robust and always outperformed those competing tools (**Figure 4B**), with the average *R*^2^ values of 0.933 (10,000 sequences), 0.947 (20,000 sequences), 0.954 (30,000 sequences), 0.960 (40,000 sequences), and 0.959 (50,000 sequences), respectively. In contrast, the corresponding average *R*^2^ values of CRPT (the second-best tool) were only 0.774 (10,000 sequences), 0.837 (20,000 sequences), 0.867 (30,000 sequences), 0.878 (40,000 sequences), and 0.889 (50,000 sequences). Next, we evaluated the performance of TRAFICA across different HT-SELEX datasets (cross-dataset experiment). For the TFs that exist in multiple HT-SELEX datasets, we extracted the fine-tuning set for TRAFICA (training set for the other tools) from one dataset and the test set from the other dataset(s). We generated 123 such datasets for 82 TFs in total (**Supplementary Table S5**). We utilized the PCC for performance assessment as we did for DREAM5 PBM datasets (**Supplementary Note 3**). TRAFICA consistently outperformed the other deep learning-based tools (**Figure 4C**), regardless of the number of sequences included in fine-tuning sets. DNAffinity and DNAshapeR are machine learning-based tools, and they can only be evaluated on 69 datasets with the TFs that have the same lengths of target sequences between training and test sets. TRAFICA still outperformed these two tools on all datasets (**Figure 4D**).

### 2.4 *In vitro* TF-DNA binding affinity prediction on DREAM5 PBM datasets using TRAFICA fine-tuned by HT-SELEX datasets

To evaluate the generalizability of TRAFICA across different assay technologies, we fine-tuned TRAFICA on HT-SELEX datasets and evaluated it on the test datasets from DREAM5 PBM datasets. We identified 86 datasets (**Supplementary Table S6**), in which the TFs both existed in HT-SELEX and DREAM5 PBM datasets. We only compared TRAFICA with those deep learning-based tools (DeepBind, DLBSS, and CRPT/CRPTS), because the lengths of target sequences of these two technologies were different (HT-SELEX: 20, 30, or 40bps; DREAM5 PBM: 35 bps). We observed that all PCC values from TRAFICA across different numbers of sequences in fine-tuning sets were better than the second-best tool DeepBind (**Figure 5A**; *p*-value=0.051 for 10,000 sequences, *p*-value=0.007 for 20,000 sequences, *p*-value=0.047 for 30,000 sequences, *p*-value=0.011 for 40,000 sequences, *p*-value=0.018 for 50,000 sequences). These results indicated that TRAFICA had better generalizability than the existing tools on datasets from different *in vitro* binding assay technologies.

### 2.5 *In vivo* TF-DNA binding affinity prediction on ChIP-seq datasets using TRAFICA fine-tuned by HT-SELEX datasets

Predicting *in vivo* TF-DNA binding affinity is more challenging than *in vitro* predictions due to the influence of multiple factors (e.g., co-binding mechanism and DNA methylation), which are dynamic in diverse cell status. Following the study of DeepBind, we fine-tuned TRAFICA on the datasets from *in vitro* HT-SELEX technology and tested the model on datasets from *in vivo* ChIP-seq technology. This strategy allows us to explore whether TRAFICA could leverage the characteristics of sequences from open chromatin regions to improve the prediction of *in vivo* TF binding affinity. This assessment was applied to 108 datasets (for the 28 TFs included in both HT-SELEX and ChIP-seq datasets, **Supplementary Table S7**), each composed of an HT-SELEX dataset and a ChIP-seq dataset. The area under the receiver operating characteristic curve (AUROC) was used as the evaluation metric, as the labels of ChIP-seq datasets were binary. TRAFICA significantly outperformed the second-best tool DeepBind in predicting *in vivo* TF-DNA binding affinity across different numbers of sequences used for model fine-tuning (training) (**Figure 5B**; *p*-value=0.007 for 10,000 sequences, *p*-value=0.008 for 20,000 sequences, *p*-value=0.010 for 30,000 sequences, *p*-value=0.011 for 40,000 sequences, *p*-value=0.049 for 50,000 sequences).

Furthermore, we performed an ablation study to evaluate the impact of the pre-training stage in predicting *in vivo* TF-DNA binding affinity. We observed the pre-training stage could improve the performance of TRAFICA with an increase of 0.13 on the average AUROC value (**Figure 5C**).

### 2.6 TF binding motif analysis

Sequence motifs are typically represented as position frequency matrices or sequence logos that provide an intuitive visualization of TF binding preferences. We used an attention-based approach (**Supplementary Note 4**, modified from DNABERT-viz [22]) to generate TF binding motifs based on the results of TRAFICA on 440 datasets (used to generate **Figure 4A**). We found 249 TFs from 362 TF-motif pairs (**Supplementary Table S8**) with the matching validated motifs in the JASPAR2022 database [23]. We employed MoSBAT energy (MoSBAT-e) scores [24] to quantify the similarity between predicted and validated motifs for the same TFs (motif pairs), and utilized “WebLogo” [25] to generate motif logos for visualization. The logos of the motif pairs with the top-3 highest similarities are shown in **Figure 6**, and their similarities are 0.8719 (TF: E2F4, JASPAR ID: MA0470.2), 0.7856 (TF: HOXA2, JASPAR ID: MA0900.2), and 0.7507 (TF: MAFK, JASPAR ID: MA0496.3), respectively. The strong agreement among these three pairs explains TRAFICA’s superior performance in predicting TF-DNA binding affinity on HT-SELEX datasets, suggesting that the model is highly effective in learning biologically relevant TF-DNA binding patterns. We showed the motif similarity distribution and motif logos for all 362 motif pairs in **Supplementary Figure S4** and **Supplementary Figure S5-S12**.

**Figure 6.**
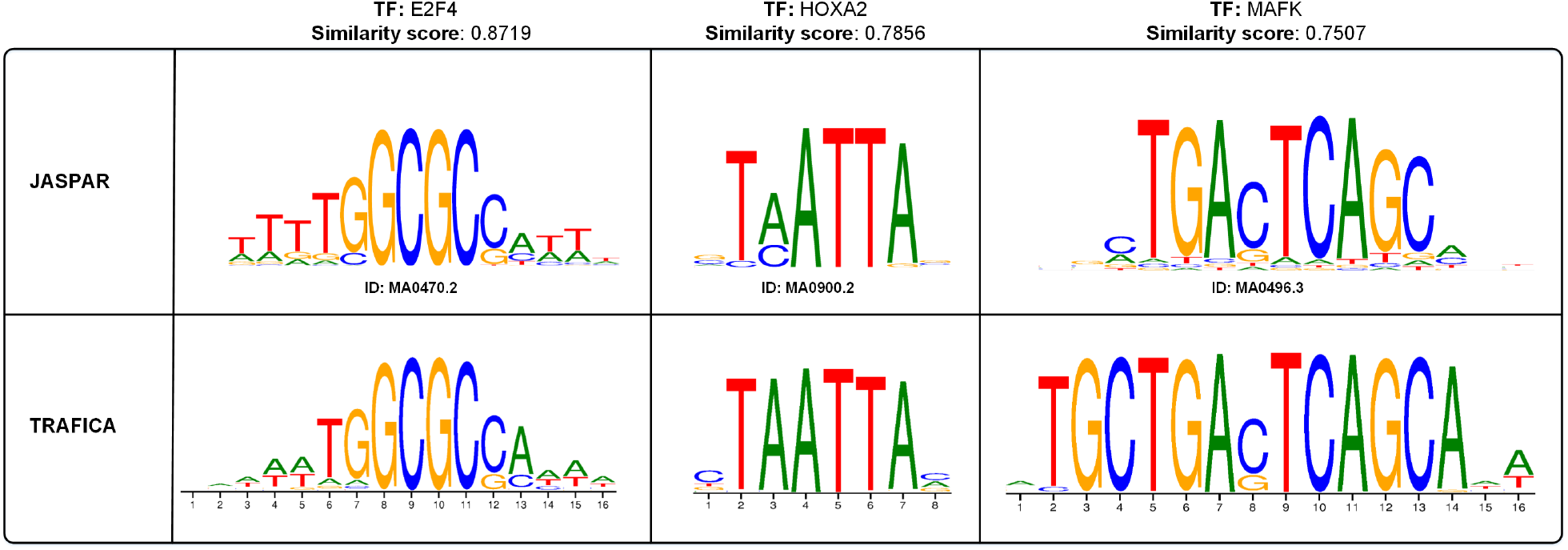
Motif logos of three TFs: E2F4, HOXA2, and MAFK. The top panel displays the validated TF binding motifs from the JASPAR2022 database, whereas the bottom panel presents the TF binding motifs generated by the attention scores from TRAFICA. The similarities between each two motifs were calculated using MoSBAT-e scores.

## 3 Discussion

The accumulation of high-throughput sequencing data on TF-DNA binding (e.g., PBM, HT-SELEX, ChIP-seq, and ATAC-seq) offers a rich landscape for delving into the binding tendencies of TFs and DNA sequences. Although ChIP-seq directly captures genome-wide DNA sequences binding to a specific TF, it has its own limitation [2] because antibodies may not always be available for all TFs. Moreover, binding patterns can vary across different cell lines for a TF [26], which can skew the understanding of intrinsic TF binding preferences. PBM technology is widely used to measure the TF-DNA binding intensities of synthesized nucleotide sequences to specific TFs by detecting fluorescence signal intensities [27]. A majority of sequences designed for PBM experiments may be with low binding affinities to the target TFs because the synthetic sequences may not closely match specific TF-binding motifs. HT-SELEX experiments can generate abundant nucleotide sequences with relatively higher binding affinities by multicycle selection (**Figure 1C**). Several studies have utilized HT-SELEX technology to investigate TF-DNA binding affinity [7, 28, 29, 30], the interactions of RNA-binding proteins [31, 32], and the interactions of drug-target proteins (e.g., aptamer-binding proteins [33]). Unlike PBM and HT-SELEX, which directly measure TF-DNA binding affinity, ATAC-seq measures chromatin accessibility that provides natural TF binding preferences. For example, promoters and enhancers are usually observed in those highly accessible (open chromatin) genomic regions [9]. We pre-trained TRAFICA using sequences from open chromatin regions annotated by the peaks from ATAC-seq experiments to enable it to learn natural TF binding preferences. Our results demonstrated that involving the pre-training stage significantly improved the performance of TRAFICA in TF-DNA binding affinity prediction. We also noticed that TRAFICA significantly outperformed DNAffinity (*p*-value<1e-3), the second-best tool, on DREAM5 PBM datasets, even without the pre-training stage. This observation could be attributed to the remarkable ability of the transformer-encoder structure in representation learning. Additionally, two pooling modules in TRAFICA can be used to generate sequence embeddings (**Figure 2B**) from token embeddings: (1) mean tokens: averaging all token embeddings out of the last transformer-encoder block; (2) [CLS] token: utilizing the embedding of “[CLS]” as the sequence embedding. We evaluated these two modules on DREAM5 PBM datasets and observed that using the mean tokens method yielded slightly better predictive performance than the [CLS] token method (**Supplementary Figure S13**). Therefore, we adopted the first module in this study.

Some studies have demonstrated DNA shape could influence protein-DNA binding in *Drosophila* [34, 35] and *S. cerevisiae* [36]. However, we observed there was no significant difference between the performance of CRPT (without DNA shape features) and CRPTS (with DNA shape features) on HT-SELEX datasets (Jolma et al.: *p*-value=0.670; Yang et al.: *p*-value=0.188; Yin et al.: *p*-value=0.437; **Figure 4A**). This suggests that DNA shape features may not consistently improve TF-DNA binding affinity prediction.

The advancement of single-cell multi-omics provides unprecedented opportunities to predict cell-type-specific TF-DNA binding affinity. Several studies [37, 38] have utilized single-cell ATAC-seq profiles to explore cell heterogeneity by analyzing cell-type-specific open chromatin regions. In addition, a previous study [39] utilized promoter sequences of target genes to predict gene expression using single-cell RNA-seq profiles, providing valuable insight into understanding gene regulation. These single-cell multi-omics technologies provide a foundation for developing tools that could potentially predict cell-type-specific TF-DNA binding affinity.

## 4 Methods

### 4.1 Data collection

We collected ATAC-seq data from the human ATAC-seq experiments in the ENCODE project [10, 11]. More specifically, we chose the experiments that were generated from the three types of biosamples: cell lines, tissues, and primary cells (defined by the ENCODE consortium), and downloaded all available “bed narrowPeak” files of these experiments (the middle panel of **Figure 1A**). We extracted the sequences from 100bps upstream and downstream flanking regions of the peaks with high confidence (−*log*_10_(*p*-value) ≥ 100, the right panel of **Figure 1A**). We utilized Bedtools [40] to retrieve the corresponding sequence from these regions in the human reference genome (GRCh38, GCF_000001405.40).

We retrieved *in vitro* TF-DNA binding profiles from the DREAM5 PBM study [19] and three previous HT-SELEX studies Jolma et al. [28] (ENA accession ID: PRJEB3289), Yang et al. [29] (ENA accession ID: PRJEB14744), and Yin et al. [30] (ENA accession ID: PRJEB9797 and PRJEB20112) to fine-tune TRAFICA, respectively. The DREAM5 PBM study comprised 132 datasets for 66 TFs (**Supplementary Table S2**), and the HT-SELEX studies included 440 datasets for 336 TFs (**Supplementary Table S3**) after rigorous quality control (**Supplementary Note 1.1**). The relative sequence binding affinities of HT-SELEX datasets were estimated based on the fold enrichment of sub-sequences between HT-SELEX selection cycles [3] (**Supplementary Note 1.2**). Moreover, we incorporated the ChIP-seq datasets from DeepBind [15] to assess the predictive capability of TRAFICA in predicting *in vivo* TF-DNA binding affinity.

### 4.2 Nucleotide sequence tokenization

We processed each nucleotide sequence into an ordered list of *k*-mer (*k* = 4 by default) tokens. In this case, each token is represented by *k* consecutive nucleotides. We constructed a vocabulary of *k*-mer tokens comprising all possible *k*-mers with four symbols (“A,” “T,” “G,” and “C”) to represent DNA nucleotides and the symbol “N” denoting unidentified nucleotides in sequences. These unidentified nucleotides may arise due to technical issues and sequencing errors. Following previous studies [20, 22], we also introduced five special tokens into the vocabulary, including “[CLS],” “[SEP],” “[MASK],” “[PAD],” and “[UNK].” The vocabulary size is 5^*k*^ + 5 (630 for *k* = 4).

### 4.3 Model pre-training

We utilized the integrated ATAC-seq dataset to pre-train TRAFICA (**Figure 1B**). Masking single tokens could oversimplify the model pre-training, as they could be completely recovered by their preceding and succeeding tokens [22]. For example, in a tokenized sequence “AGAA [MASK] AACG,” the masked token can be ascertained as “GAAC” by its neighbor tokens. To address this issue, we designed a strategy (**Supplementary Note 5**) to mask consecutive tokens (e.g., “AGAA [MASK] [MASK] ACGT CGTA” or “AGAA GAAC [MASK] [MASK] CGTA”) instead of masking single tokens for model pre-training. In this study, the proportion of masked tokens in each sequence is 15% (by default). Given a sequence with masked tokens {*t*_1_, *t*_2_, …, *t*_*M*_}, we calculated the cross-entropy loss as follows:

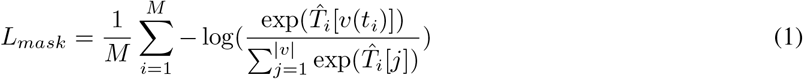

where *M* represents the number of masked tokens in the sequence. *v*(·) denotes the index of a specific token in the token vocabulary, and |*v*| represents the size of the token vocabulary. 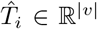 is the output of the token predictor for *i*-th masked token (**Figure 2A**). Given an ATAC-seq dataset (denoted as *D*_0_), we minimized the cross-entropy of masked tokens in the pre-training stage:

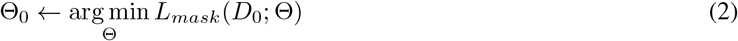

where Θ and Θ_0_ represent the initial and pre-trained model parameters, respectively.

TRAFICA was pre-trained over a total number of 110,000 steps using a batch size of 384. We employed a linear warm-up strategy to select the learning rates during the pre-training stage (from 0 to 0.0001 over the first 10,000 steps). Subsequently, we decayed the learning rate to 0 over the remaining steps. We applied the AdamW optimizer [41] with the hyperparameters *β*_1_ = 0.9, *β*_2_ = 0.98, and *ϵ* = 10^−7^ to calculate the gradient and optimize the model parameters. The pre-training process was executed on an Nvidia Tesla A100 GPU card.

### 4.4 Model fine-tuning

TRAFICA was fine-tuned on the TF-DNA binding profiles from PBM and HT-SELEX experiments (**Figure 1C** and **D**). Given a specific TF-DNA binding dataset (denoted as *D′*) with *N* sequences and corresponding affinities {*y*_1_, *y*_2_, …, *y*_*N*_}, we minimized the mean squared error (MSE) as follows:

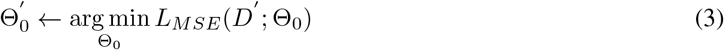

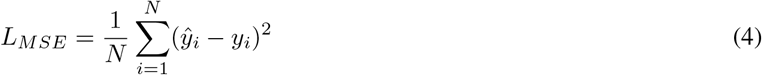

where 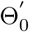 represents the parameters of the model fine-tuned on *D*′. *ŷ*_*i*_ is the output of the affinity predictor for *i*-th sequence (**Figure 2B**). We employed 10% of the fine-tuning dataset as a validation set for early stopping. The remaining 90% was utilized to optimize the model parameters using the AdamW optimizer with a constant learning rate of 0.00002 and a batch size of 128. Besides, the training settings for the comparison tools used in this study are included in **Supplementary Note 6**.

### 4.5 HT-SELEX subset construction

We generated subsets by sampling different numbers of sequences from HT-SELEX datasets to fine-tune TRAFICA, evaluating the impact of the number of sequences used for model fine-tuning. We applied a binning technique to a target fine-tuning dataset to group sequences based on their binding affinities, resulting in *N*_*sub*_ bins. Next, we randomly selected a sequence from each bin to construct a subset with *N*_*sub*_ sequences. More specifically, denoted *v*_*max*_ (*v*_*min*_) as the maximum (minimum) affinity value, we calculated the range of bins as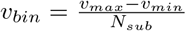. Thus, the edges of bins are {[*v*_*min*_, *v*_*min*_ + *v*_*bin*_), [*v*_*min*_ + *v*_*bin*_, *v*_*min*_ + 2 *× v*_*bin*_), …, [*v*_*min*_ + (*N*_*sub*_ − 1) *× v*_*bin*_, *v*_*max*_]}.

## Data availability

The DREAM5 PBM experimental data is available at https://hugheslab.ccbr.utoronto.ca/supplementary-data/DREAM5/. The HT-SELEX experimental data can be accessed by the ENA accession ID: PRJEB3289 (Jolma et al.), PRJEB14744 (Yang et al.), and PRJEB9797/PRJEB20112 (Yin et al.). The code of TRAFICA is available at https://github.com/ericcombiolab/TRAFICA. The pre-trained model is available at https://huggingface.co/Allanxu/TRAFICA. The processed sequencing data of ATAC-seq, HT-SELEX, PBM, and ChIP-seq can be accessed from https://zenodo.org/doi/10.5281/zenodo.8248339.

## Author contributions statement

LZ and YX conceived the study; YX designed TRAFICA; YX implemented the algorithm; YX conducted the experiments and analyzed the results; LZ and YX wrote the article; CHW, KX, YD, and APL reviewed the paper. All authors read and approved the final manuscript.

## Acknowledgments

This research is partially supported by Young Collaborative Research Grant (C2004-23Y), Health and Medical Research Fund (11221026), HKBU Start-up Grant Tier 2 (RC-SGT2/19-20/SCI/007).

