## Supplementary Notes/Figures for "TRAFICA: An Open Chromatin Language Model to Improve Transcription Factor Binding Affinity Prediction"

### Supplementary File for *TRAFICA: An Open Chromatin Language Model to Improve Transcription Factor Binding Affinity Prediction*

Yu Xu[1], Chonghao Wang[1], Ke Xu[1], Yi Ding[1], Aiping Lyu[2,*], and Lu Zhang[1,3,*]

[1]Department of Computer Science, Hong Kong Baptist University
[2]School of Chinese Medicine, Hong Kong Baptist University
[3]Institute for Research and Continuing Education, Hong Kong Baptist University

### Supplementary Notes

#### Supplementary Note 1: HT-SELEX data source

We downloaded the following HT-SELEX experimental data files from the ENA [1]: 1. Jolma et al. [2]: 547 HT-SELEX experiments covering 461 unique human and mouse TFs; 2. Yang et al. [3]: 521 HT-SELEX experiments covering 445 unique human and mouse TFs; 3. Yin et al. [4]: 2265 HT-SELEX experiments covering 550 unique human TFs.

##### 1.1: Quality control for HT-SELEX experiments

We filtered out the experiments that lacked the following information: Ensembl ID, structural domains, and amino acid sequences. To ensure the reliability of HT-SELEX experiments, we filtered out low-quality experiments by conducting quality control. Given an HT-SELEX experiment with $N$ selection cycles, we implemented quality control as follows:

1. For $i$-th ($i \neq 0$) selection cycle, we counted the frequency of all possible 8-mers, denoting as $F_i = \{f_i^1, ..., f_i^{65536}\}$ ($65536 = 4^8$: the number of all 8-mers).
2. For the initial cycle, we estimated the expected frequency of all possible 8-mers by using a fifth-order Markov model, denoting as $E_0 = \{f_0^1, ..., f_0^{65536}\}$. The Markov model was implemented using the R programming package "SELEX" [5].
3. Following the previous study [6], we defined the ratio of $F_i$ to $E_0$ as the enrichment score of 8-mers in the $i$-th cycle, denoting as $ES_i = \{\frac{f_i^1}{f_0^1}, ..., \frac{f_i^{65536}}{f_0^{65536}}\}$).
4. Following the previous studies [7, 8], we excluded the experiment if it satisfies one of the following criteria: (a) The Spearman's rank correlation coefficient of the enrichment scores $ES_3$ and $ES_4$ is less than 0.8, which implies the neighboring cycles are uncorrelated in the enrichment patterns; (b) The number of sequences with counts larger than 10 in the $N$-th (final) selection cycle is less than 500; (c) The count of each possible 8-mer in the initial cycle is less than 100, which could result in inaccurate estimates of $E_0$.

After implementing the above steps, 440 high-quality experiments remained, consisting of 336 unique TFs.

##### 1.2: Relative binding affinity estimation for HT-SELEX data

For each high-quality HT-SELEX experiment, we utilized the R programming package "SELEX" [5] to estimate relative $k$-mer binding affinities. We employed a searching range of $\{3, 4, ..., 13, 14\}$ for the optimal hyper-parameter $k$, as the minimum sequence length of the HT-SELEX datasets used in this study is 14. We then merged the estimated $k$-mer binding affinities for each sequence to obtain sequence affinities. Given a sequence $a$, we merge the $k$-mer binding affinities $A_{kmer}$ into the sequence affinity $A_{seq}(a)$ by the following:

$$A_{seq}(a) = \frac{\sum_{i=1}^{K} A_{kmer_i}}{\sqrt{K}} \quad (1)$$

where $K$ represents the number of $k$-mers in the sequence $a$. This approach (Eq. 1) was introduced in the previous study [9] that integrated the statistics of $k$-mers to generate relative sequence binding affinities. Additionally, we removed the low-complexity sequences that present a DUST [10] score larger than 2, as low-complexity sequences may consist of repeated $k$-mers with a high relative k-mer binding affinity. This could cause inaccurate sequence binding affinity. Given a sequence $a$ with a length $n > 2$, its DUST score $S(a)$ is formulated as follows:

$$S(a) = \frac{\sum_{t \in R} c_t(a)(c_t(a) - 1)/2}{l - 1} \tag{2}$$

where $R$ is a set of 64 possible triplets of DNA base combinations (e.g., AAA, AAT, ATT,...). $l = n - 2$ is the number of triplets in the sequence $a$. $c_t(a)$ represents the frequency of occurrence of the triplet $t$ within the sequence $a$.

The sequences and corresponding binding affinities from the final selection cycles are used for model fine-tuning, as the final cycle typically contains a higher ratio of high-affinity sequences than the early cycles. Furthermore, we noticed that the HT-SELEX experiments conducted by Yang et al. [3] have a notably higher read depth compared to Jolma et al [2], with an almost 10-fold increase in the average number of reads per sequencing file (from 168,000 to 1,656,000). Those HT-SELEX experiments (Yang et al.) generated large numbers of sequences, which significantly increases the time spent on the fine-tuning stage and could potentially destroy the knowledge learned during the pre-training stage. To alleviate this issue, we limited our analysis to the first 0.5 million sequences ordered by relative binding affinities for each dataset from Yang et al. [3], as we observed that most datasets from the three studies contained fewer than 0.5 million sequences (**Figure S14**).

#### Supplementary Note 2: Model structure of TRAFICA

##### 2.1: Embedding layers

The token embedding and position embedding layers are two fully connected networks, denoted as $W_t \in \mathbb{R}^{d_{vocab} \times d}$ and $W_p \in \mathbb{R}^{l_{max} \times d}$, where $d_{vocab}$ and $l_{max}$ represent the size of the token vocabulary and the maximum number of input tokens. Given a sequence with a length $l_s \geq 4$, it is tokenized as a list of 4-mer tokens and two specific tokens '[CLS]' and '[SEP]' at both ends. The length of this token list is $l = l_s - 3 + 2$. Each token $t$ and the corresponding position index $p$ are converted as two one-hot encoding vectors $x_t \in \mathbb{R}^{d_{vocab}}$ and $x_p \in \mathbb{R}^{l_{max}}$, respectively. The process of the embedding layers can be formulated as follows:

$$h_t = x_t^T W_t \oplus x_p^T W_p \tag{3}$$

$$H^0 = Concat(h_1, h_2, ...h_l) \tag{4}$$

where $h_i \in \mathbb{R}^d$ is the hidden embedding capturing the semantic and positional information of the token $t_i$, and $d$ denotes the dimension of the hidden embedding. $Concat(\cdot)$ is the operation of concatenation. The matrix $H^0 \in \mathbb{R}^{l \times d}$ of the sequence is the input for the first transformer-encoder block.

##### 2.2: Transformer-encoder

The structure of the transformer-encoder is composed of four components, a multi-head self-attention module, a feed-forward module, layer normalization, and residual connections. Each transformer-encoder block takes as input the output from the previous block, except the first block. Given the transformer-encoder block with $n_{head}$ attention heads, the dimension of hidden representation $d_k = d/n_{head}$ in each head, and an input matrix $X \in \mathbb{R}^{l\times d}$, the process of the multi-head self-attention module can be described as follows:

$$MultiHead(X) = Concat(AttHead_1, AttHead_2, ...AttHead_{n_{head}})W^O \tag{5}$$

where $W^O \in \mathbb{R}^{d\times d}$ is the learnable parameters used to aggregate the outputs of attention heads. The self-attention mechanism for each head is defined as follows:

$$AttHead_i = Softmax(\frac{XW_i^Q \otimes (XW_i^K)^T}{\sqrt{d_k}})XW_i^V \tag{6}$$

where $\{W_i^Q \in \mathbb{R}^{d\times d_k}, W_i^K \in \mathbb{R}^{d\times d_k}, W_i^V \in \mathbb{R}^{d\times d_k}\}_{i=1}^{n_{head}}$ are the learnable parameters of the linear projection for each attention head, and $\sqrt{d_k}$ is the scaling factor. $\otimes$ denotes the operation of dot-production. Given an input $X \in \mathbb{R}^{l\times d}$, the feed-forward module is formulated as follows:

$$FNN(X) = GELU(XW_1^{FNN} + b_1^{FNN})W_2^{FNN} + b_2^{FNN} \tag{7}$$

where $\{W_1^{FNN} \in \mathbb{R}^{d\times d_{FNN}}, b_1^{FNN} \in \mathbb{R}^{d_{FNN}}, W_2^{FNN} \in \mathbb{R}^{d_{FNN}\times d}, b_2^{FNN} \in \mathbb{R}^d\}$ are the learnable parameters of the first and second fully connected layers in this module. $d_{FNN}$ represents the dimension of immediate embeddings in the feed-forward module. $GELU(\cdot)$ is the non-linear activation function. Based on **Equation 5** and **7**, the forward propagation of the transformer-encoder blocks is depicted as follows:

$$H_1 = LayerNorm(MultiHead(H) \oplus H) \tag{8}$$

$$H_2 = LayerNorm(FNN(H_1) \oplus H_1) \tag{9}$$

where $H \in \mathbb{R}^{l\times d}$ represents the output from the previous block ($H = H^0$ for the first block and $H = H_2^{i-1}$ for the $i$-th block). $\oplus$ denotes the operation of the element-wise addition.

##### 2.3: Token predictor

The token predictor consists of two fully connected layers with a GELU activation function and layer normalization for performing masked token prediction in the pre-training phase. Given the output of the last transformer-encoder block $H^N$, the operation of the token predictor is defined as follows:

$$\hat{T} = LayerNorm(GELU(H^{n_{blocks}}W_1^t + b_1^t))W_2^t + b_2^t \tag{10}$$

where $\{W_1^t \in \mathbb{R}^{d\times d}, b_1^t \in \mathbb{R}^d, W_2^t \in \mathbb{R}^{d\times|v|}, b_2^t \in \mathbb{R}^{|v|}\}$ are the parameters of the predictor. $H^{n_{blocks}} \in \mathbb{R}^{l\times d}$ represents the output of the last transformer block. The predicted scores are denoted as $\hat{T} \in \mathbb{R}^{l\times|v|}$, where $|v|$ is the size of the token vocabulary.

##### 2.4: Pooler module

The pooler module is composed of a fully connected layer with a Tanh activation function to transform the token embeddings $H^{n_{blocks}} = \{h_1^{n_{blocks}}, h_2^{n_{blocks}}, ...h_l^{n_{blocks}}\}$ into a sequence embedding $h_{seq} \in \mathbb{R}^{1\times d}$. The process of obtaining sequence embeddings is formulated as follows:

$$h_{seq} = Tanh(h_{mean}^{n_{blocks}} W^{pool} + b^{pool}) \tag{11}$$

where

$$h_{mean}^{n_{blocks}} = \frac{1}{l-2} Concat(h_2^{n_{blocks}}, h_3^{n_{blocks}}, ..., h_{l-1}^{n_{blocks}}) \tag{12}$$

where $\{W^{pool} \in \mathbb{R}^{d\times d}, b^{pool} \in \mathbb{R}^d\}$ is the parameters of the pooler module. We remove the embeddings of the tokens '[CLS]' and '[SEP]' in generating the sequence embedding.

##### 2.5: Affinity predictor

We employed two fully connected layers as a regression predictor to transform the sequence embedding $h_{seq}$ into a scalar value of the predicted affinity $\hat{y}$. The process of this predictor is shown as follows:

$$\hat{y} = (h_{seq} W_1^{reg} + b_1^{reg}) W_2^{reg} + b_2^{reg} \tag{13}$$

where $\{W_1^{reg} \in \mathbb{R}^{d\times\lfloor d/2\rfloor}, b_1^{reg} \in \mathbb{R}^{\lfloor d/2\rfloor}, W_2^{reg} \in \mathbb{R}^{\lfloor d/2\rfloor\times 1}, b_2^{reg} \in \mathbb{R}^1\}$ are the parameters of the predictor.

#### Supplementary Note 3: Evaluation metrics

Pearson correlation coefficient (PCC) and coefficient of determination ($R^2$) are utilized to evaluate the performance in predicting *in vitro* TF-DNA binding affinity. Given a set of input sequences $X = \{x_1, x_2, ..., x_n\}$ with the label $Y = \{y_1, y_2, ..., y_n\}$ and the predicted affinities $f(X) = \{f(x_1), f(x_2), ..., f(x_n)\}$, the computation of these two metrics is formulated as follows:

$$PCC(Y, f(X)) = \frac{\sum_{i=1}^n((y_i - \bar{y})(f(x_i) - \overline{f(x)}))}{\sqrt{\sum_{i=1}^n(y_i - \bar{y})^2 \sum_{i=1}^n(f(x_i) - \overline{f(x)})^2}} \tag{14}$$

$$R^2(Y, f(X)) = 1 - (\frac{\sum_{i=1}^n(y_i - f(x_i))^2}{\sum_{i=1}^n(y_i - \bar{y})^2}) \tag{15}$$

where $f$ and $n$ represent the prediction model and the number of sequences in this set, respectively. $\overline{f(x)} = \frac{1}{n}\sum_{i=1}^n f(x_i)$ is the mean value of predicted affinities.

We employed PCC as the evaluation metric instead of $R^2$ in the following scenarios: 1. the prediction on PBM datasets; 2. the cross-datasets evaluation (fine-tuning on an HT-SELEX dataset and testing on other HT-SELEX datasets); and 3. the cross-technologies evaluation (fine-tuning on HT-SELEX datasets and testing on PBM datasets). This choice was made due to the incomparable values of labels between the datasets used for fine-tuning and testing, which could result in an $R^2$ value approaching zero or even turning negative. For instance, TRAFICA achieved a PCC of 0.807 in predicting binding affinity for TF *Nxk2* but resulted in a negative value for $R^2$. We plotted the relative affinity distributions of TF *Nxk2*, and observed that the binding affinity distributions of the fine-tuning (training for other tools) set and the predicted set show similar shapes, while differing from the binding affinity distribution of the test set (**Supplementary Figure S3**). Therefore, the $R^2$ value computed between the test set and the predicted set, whose affinity values are not directly comparable, is negative. Additionally, AUROC is utilized to evaluate the performance of predicting *in vivo* TF-DNA binding affinity on ChIP-seq datasets, considering the binary nature of labels in ChIP-seq datasets.

**Supplementary Note 4: TF binding motif analysis based on attention scores**

We employed a modified version of DNABERT-viz [11] to derive TF binding motifs based on the attention scores computed using high-affinity sequences (top-1000) from HT-SELEX datasets through TRAFICA. The attention scores of the tokens in an input sequence can be computed as follows: (1) In each attention head, we calculated a self-attention matrix through the dot-production of Query and Key embeddings in the multi-head self-attention module (**Supplementary Note 2.2**), and discard the elements corresponding "[CLS]" and "[SEP]" tokens in the self-attention matrix. (2) We then summarized the self-attention matrix by rows to obtain the attention vector representing the average attention of each token received from other tokens (including itself). (3) As we applied 12 transformer-encoder blocks and 12 attention heads in each block, we thus had $12 \times 12$ self-attention matrices for an input sequence. We derived the attention vectors from these self-attention matrices by conducting steps 1 and 2 above. This resulted in $12 \times 12$ attention vectors for the input sequence, which were then collected and averaged into one attention vector.

Given an input sequence with $l$ tokens, we defined high-attention tokens as those with an attention score larger than the mean score (averaging the attention scores of all tokens). Subsequently, we identified sub-sequences with a minimum number ($l_{min} = 4$) of contiguous high-attention tokens. We treated these sub-sequences as motif candidates and recorded their positions in the input sequence (denoted the position of a motif candidate as $[p_{start}, p_{end}]$; $l$>($p_{end}$-$p_{start}$)). We retrieved motif candidates from top-1000 high-affinity sequences and derived TF binding motifs from these candidates as follows: (1) Most motif candidates obtained from HT-SELEX sequences show considerable resemblance with some being identical, which can be merged. We performed pairwise alignment to merge a set of similar motif candidates as a motif instance by utilizing Biopython [12], where each motif instance can eventually be converted into a TF binding motif. More specifically, we arranged the list of motif candidates in descending order based on their length ($A = \{motif_1, motif_2, ...motif_n\}$, $l_{motif_1} \geq l_{motif_2}$). We utilized a separate list $B$ to accumulate motif instances and included the first element in list $A$ into list $B$ as the first motif instance ( $A = \{motif_2, ...motif_n\}$,

$B = \{motif_1\}$). Next, for each $motif_i \in A$, we tried to align it with $motif_j \in B$ to determine whether they can be merged. We ascertained that the alignment of $motif_i$ and $motif_j$ is successful if their alignment score exceeded $\max(l_{min} - 1, 0.5 \times \min(l_{motif_i}, l_{motif_j}))$, where the alignment score refers to the number of matched nucleotides between them. If $motif_i$ can be aligned with $motif_j$, we updated the position of $motif_i$ ($[p_{start}^{motif_i}, p_{end}^{motif_i}]$) by calculating positional offsets according to the alignment shift, insertions, and deletions in the alignment result. We then added $motif_i$ with its updated position into a sub-list $B_{motif_j}$ that contains the motif candidates merged with the motif instance $motif_j$. If $motif_i$ cannot be aligned with $motif_j$, we added $motif_i$ into list $B$ as a new motif instance, allowing $motif_i$ to be aligned with other motif candidates of list $A$ in the next iteration. (2) For the motif instance $motif_{max} \in B$ with the maximum number of merged motif candidates, we used the updated positions of the merged motif candidates $motif_k \in B_{motif_{max}}$ to match the regions of their corresponding sequences (top-1000 sequences). We extracted a sub-sequence with a length of $l_{sub}$ ($l_{sub}$=8 for 20-bps sequences; $l_{sub}$=16 for 30 and 40-bps sequences) from each matched region. These extracted sub-sequences were then used to construct a PFM representing a TF binding motif.

**Supplementary Note 5: Token masking for TRAFICA pre-training**

We proposed a masking strategy for TRAFICA pre-training, which masks multiple distinct regions of consecutive tokens in a nucleotide sequence. Considering a sequence with $n_t$ ($n_t = 197$ for genomic sequences used in the pre-training stage) tokens and a masked ratio of 15%, the number of masked tokens is $n_m = \lceil 0.15 \times n_t \rceil$. We randomly masked multiple regions within a sequence, with each region covering $n_c$ consecutive tokens. Thus, the number of the masked regions can be calculated as $n_r = n_m / n_c$. We selected the model employing this masking strategy with $n_c = 3$ ($n_r = (\lceil 0.15 \times n_t \rceil)/n_c = 10$) during the pre-training stage, as it achieved the highest pre-training accuracy of approximately 80% throughout the pre-training stage (the right panel of **Figure S15**).

**Supplementary Note 6: Training settings for comparison tools**

We followed the default training setting of CRPTS [13] to train deep learning-based tools (DeepBind, DLBSS, CRPT, and CRPTS), using an Adadelta optimizer with a learning rate of 0.01, a batch size of 300, and the maximum 100 epochs. The parameters of the Adadelta optimizer and the dropout rate in neural network layers were sampled within the pre-defined range (Delta:{0.9, 0.99, 0.999}; Moment: {1e-04, 1e-06, 1e-08}; Dropout:{0.2,0.5}). The training procedure for each dataset was repeated 20 times to select the optimal parameters of the Adadelta optimizer and the dropout rate. For training on HT-SELEX subsets, we adjusted the learning rate to 0.1, as we observed that the models could not be optimal during 100 epochs using the default setting. We utilized the R programming package "DNAshapeR" to extract the DNA shape features from sequences, and used the Python programming package "scikit-learn" [14] ("sklearn.ensemble.RandomForestRegressor" with the default parameters) to implement the random forest model for TF-DNA binding affinity prediction. Regarding the implementation of DNAffinity, we applied its default setting to train and test models.

#### Supplementary Tables

- Supplementary Table S1: Details of the collected ENCODE ATAC-seq experiments
- Supplementary Table S2: Details of the collected PBM experiments
- Supplementary Table S3: Details of the collected HT-SELEX experiments
- Supplementary Table S4: Model configuration of TRAFICA
- Supplementary Table S5: Datasets for HT-SELEX cross-dataset evaluation
- Supplementary Table S6: Datasets for *in vitro* cross-technologies evaluation
- Supplementary Table S7: Datasets for *in vivo* cross-technologies evaluation
- Supplementary Table S8: Motif comparisons

#### Supplementary Figures

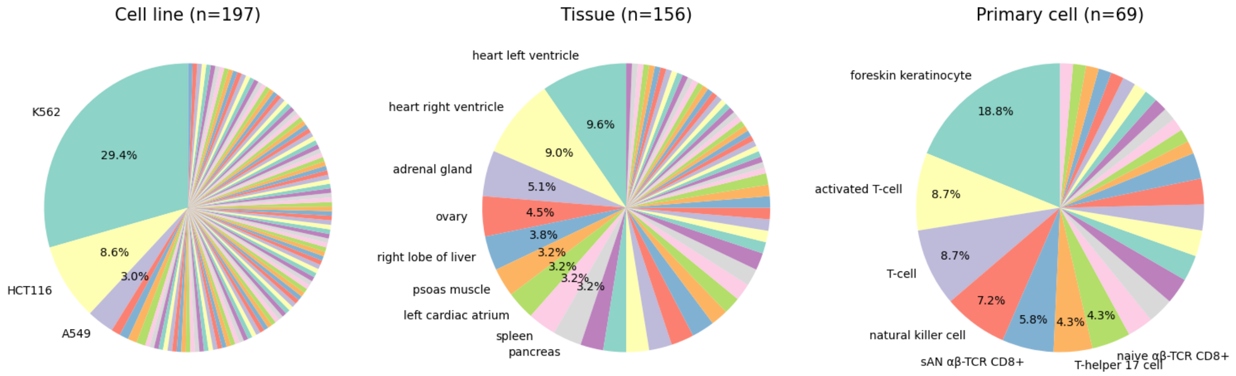


**Figure S1.** The provenance of ATAC-seq data. A total of 422 human ATAC-seq experimental profiles from the ENCODE project were collected, comprising 197 from distinct cell lines, 156 from various tissues, and 69 from diverse primary cells, respectively. The pie plots do not indicate the biosample types with a proportion of less than 3%. Abbreviations: sAN $\alpha\beta$-TCR CD8+ (stimulated activated naive CD8-positive, alpha-beta T cell), naive $\alpha\beta$-TCR CD8+ (naive thymus-derived CD8-positive, alpha-beta T cell).

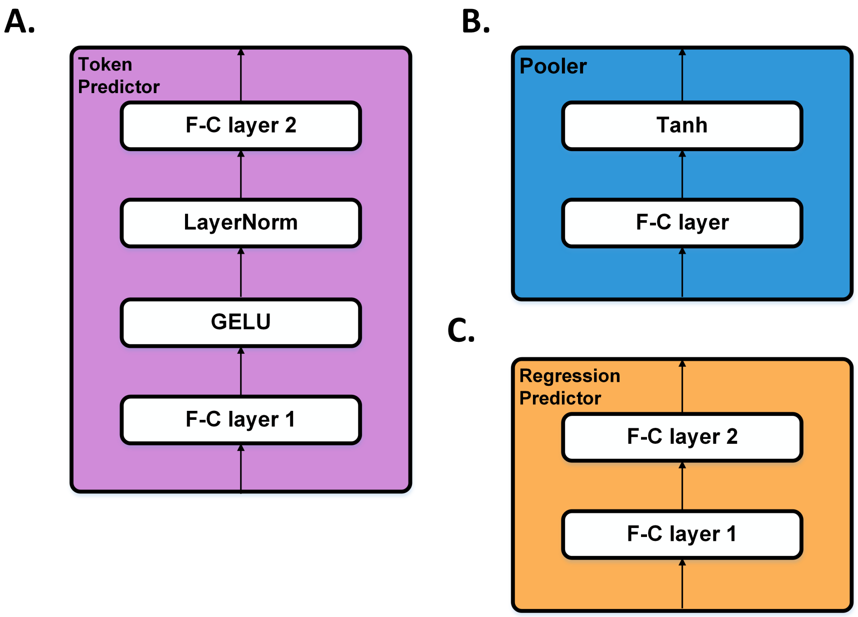


**Figure S2.** The modules of token predictor, pooler, and regression (affinity) predictor. **(A)** Token predictor consists of 4 sub-components, including two fully connected (F-C) layers ($\{W_1^t \in \mathbb{R}^{d\times d}$,$b_1^t \in \mathbb{R}^d\}$, $\{W_2^t \in \mathbb{R}^{d\times|v|}, b_2^t \in \mathbb{R}^{|v|}\}$), a GELU activation, and LayerNorm. **(B)** Pooler module comprises an F-C layer ($\{w^{pool} \in \mathbb{R}^{d\times d}, b^{pool} \in \mathbb{R}^d\}$) and a nonlinear activation function Tanh. **(C)** Regression (affinity) predictor comprises two F-C layers ($\{W_1^{reg} \in \mathbb{R}^{d\times\lfloor d/2\rfloor}, b_1^{reg} \in \mathbb{R}^{\lfloor d/2\rfloor}, W_2^{reg} \in \mathbb{R}^{\lfloor d/2\rfloor\times 1}, b_2^{reg} \in \mathbb{R}^1\}$). $d$ and $|v|$ represent the dimension of token embeddings and the size of the token vocabulary, respectively.

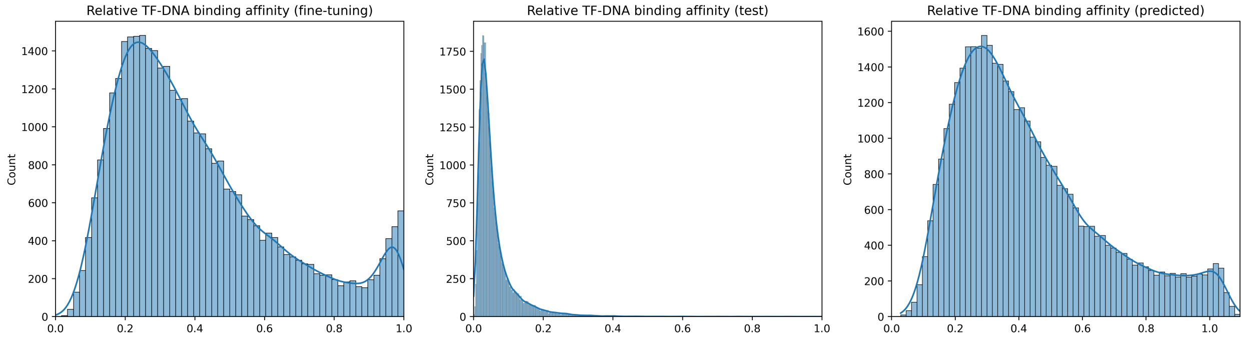


**Figure S3.** Relative affinity distribution on the PBM datasets for TF Nkx2. (**Left panel**) The TF-DNA binding affinities of sequences used for model fine-tuning. (**Middle panel**) The TF-DNA binding affinities of sequences used for model testing. (**Right panel**) The TF-DNA binding affinities of testing sequences predicted by TRAFICA.

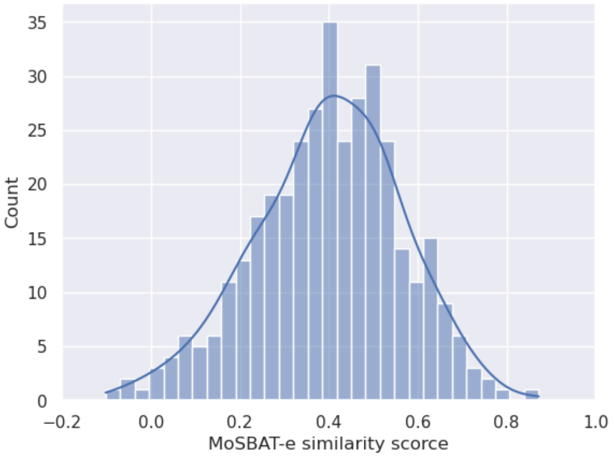


**Figure S4.** The distribution of MoSBAT-e similarity scores for 362 motif pairs.

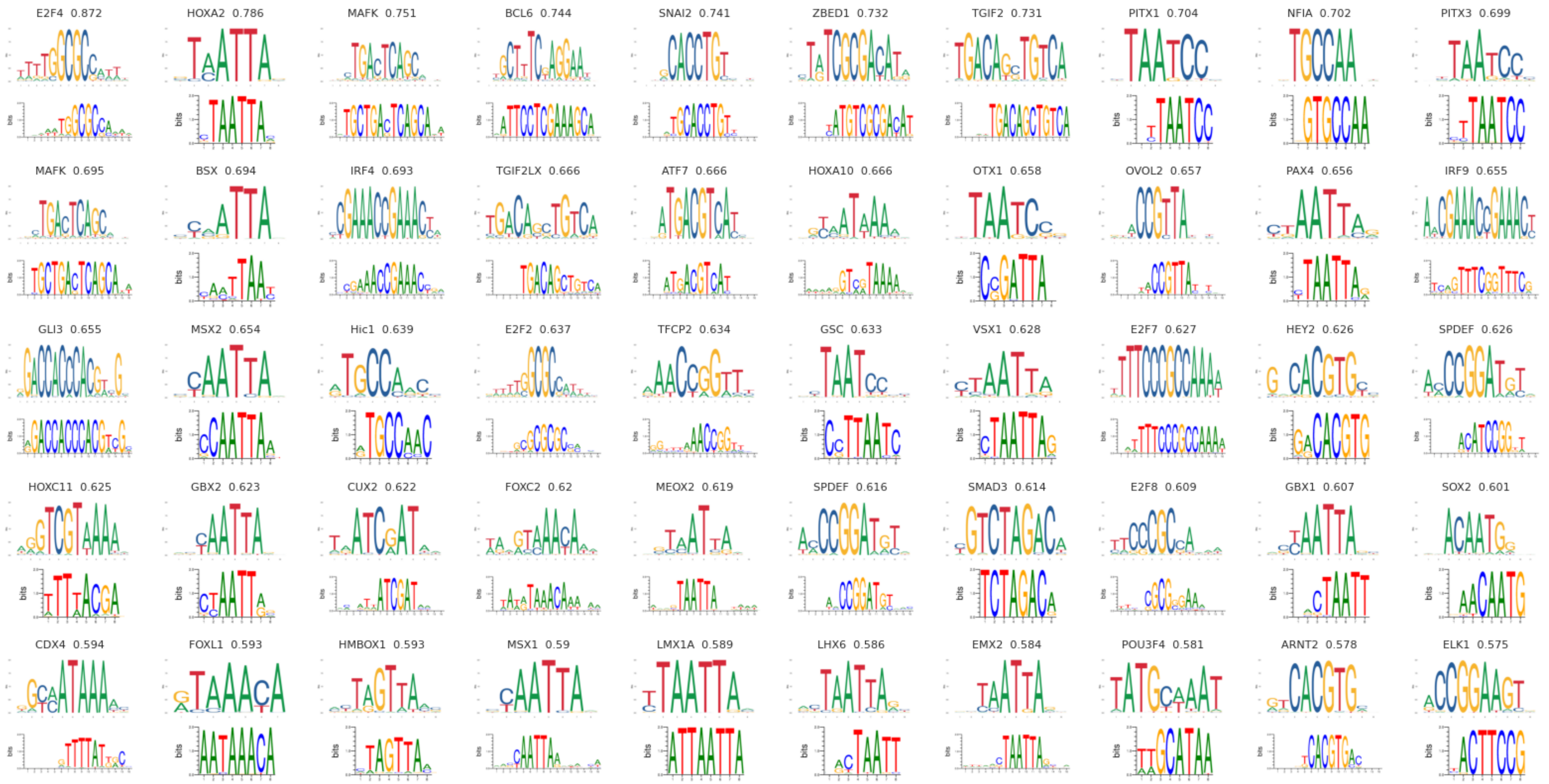


**Figure S5.** The logos of the motif pairs with the top-50 highest similarities. The top panel displays the validated TF binding motifs from the JASPAR2022 database, whereas the bottom panel presents the TF binding motifs generated by the attention scores from TRAFICA. The titles of each subfigure indicate TF names and similarity scores.

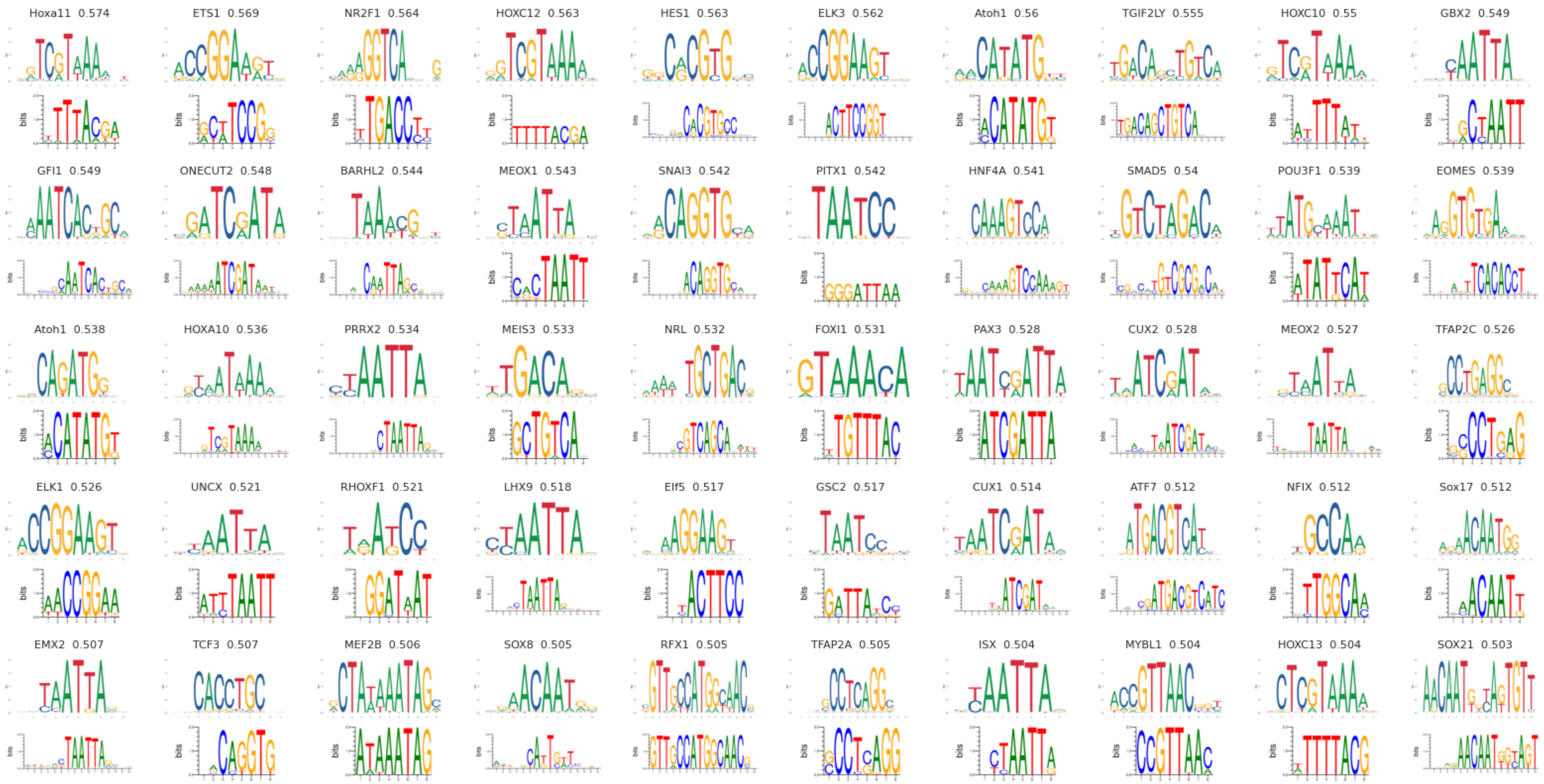


**Figure S6.** The logos of the motif pairs whose similarities ranked from 51 to 100.

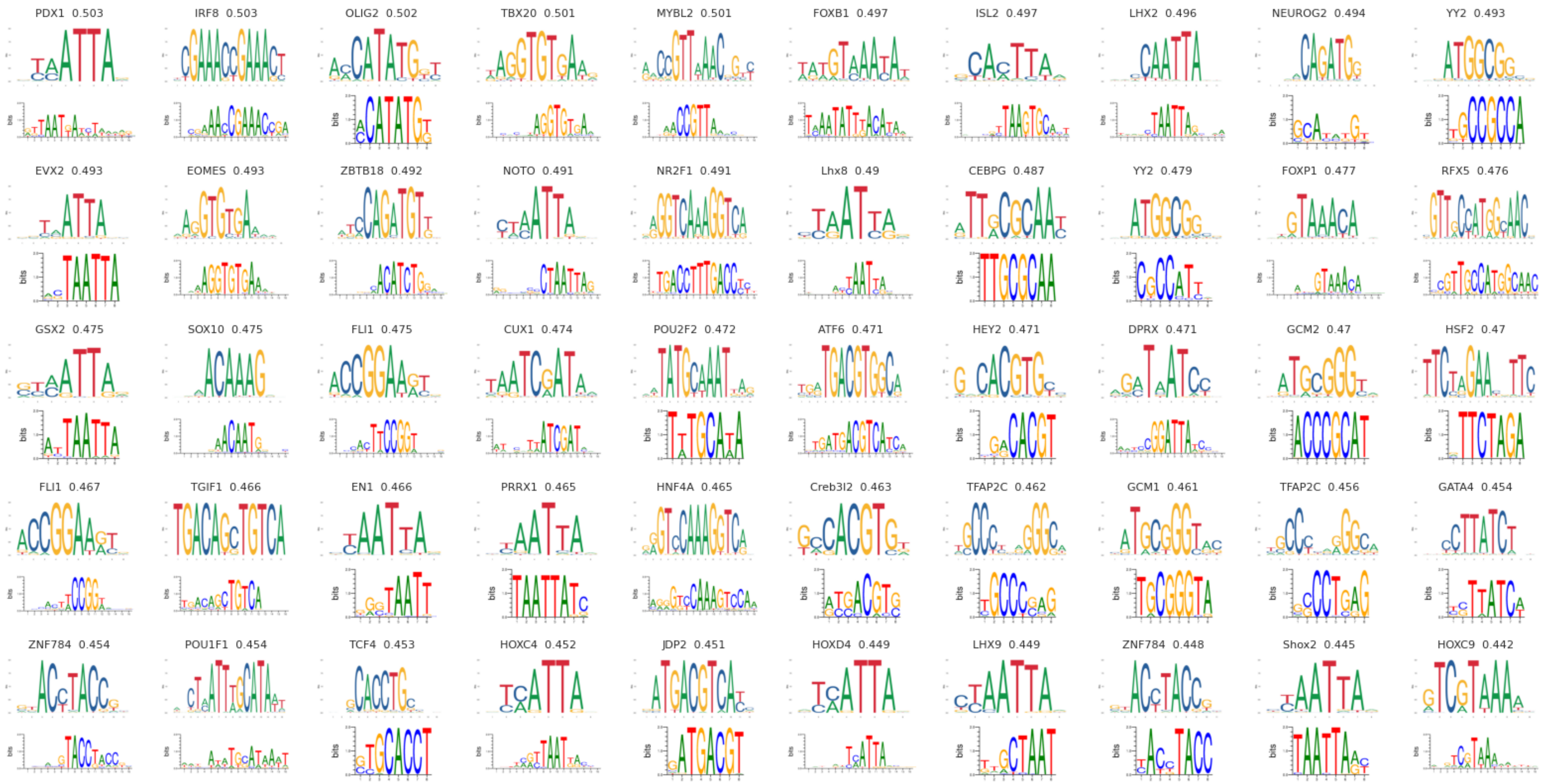


**Figure S7.** The logos of the motif pairs whose similarities ranked from 101 to 151.

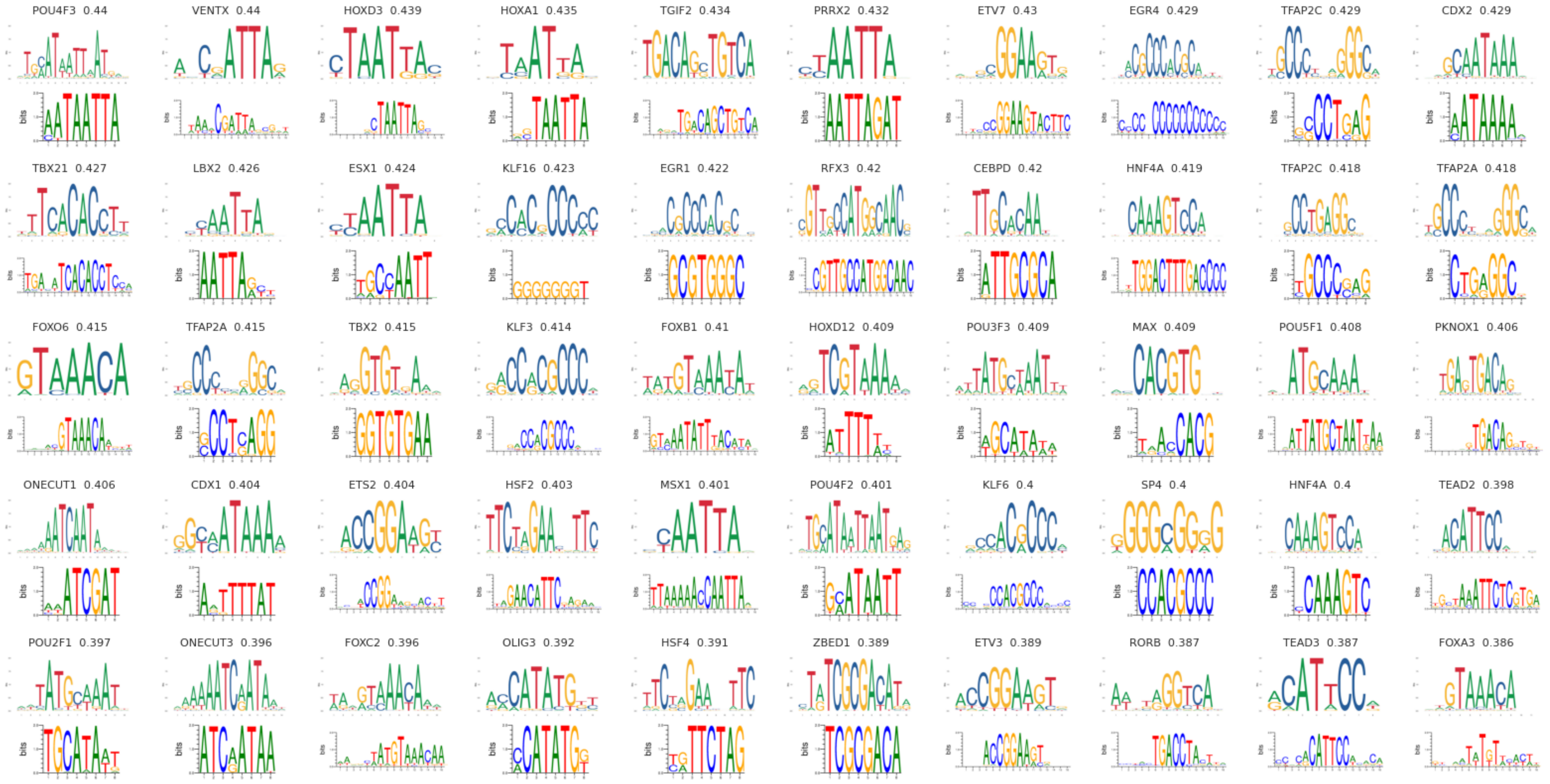


**Figure S8.** The logos of the motif pairs whose similarities ranked from 151 to 200.

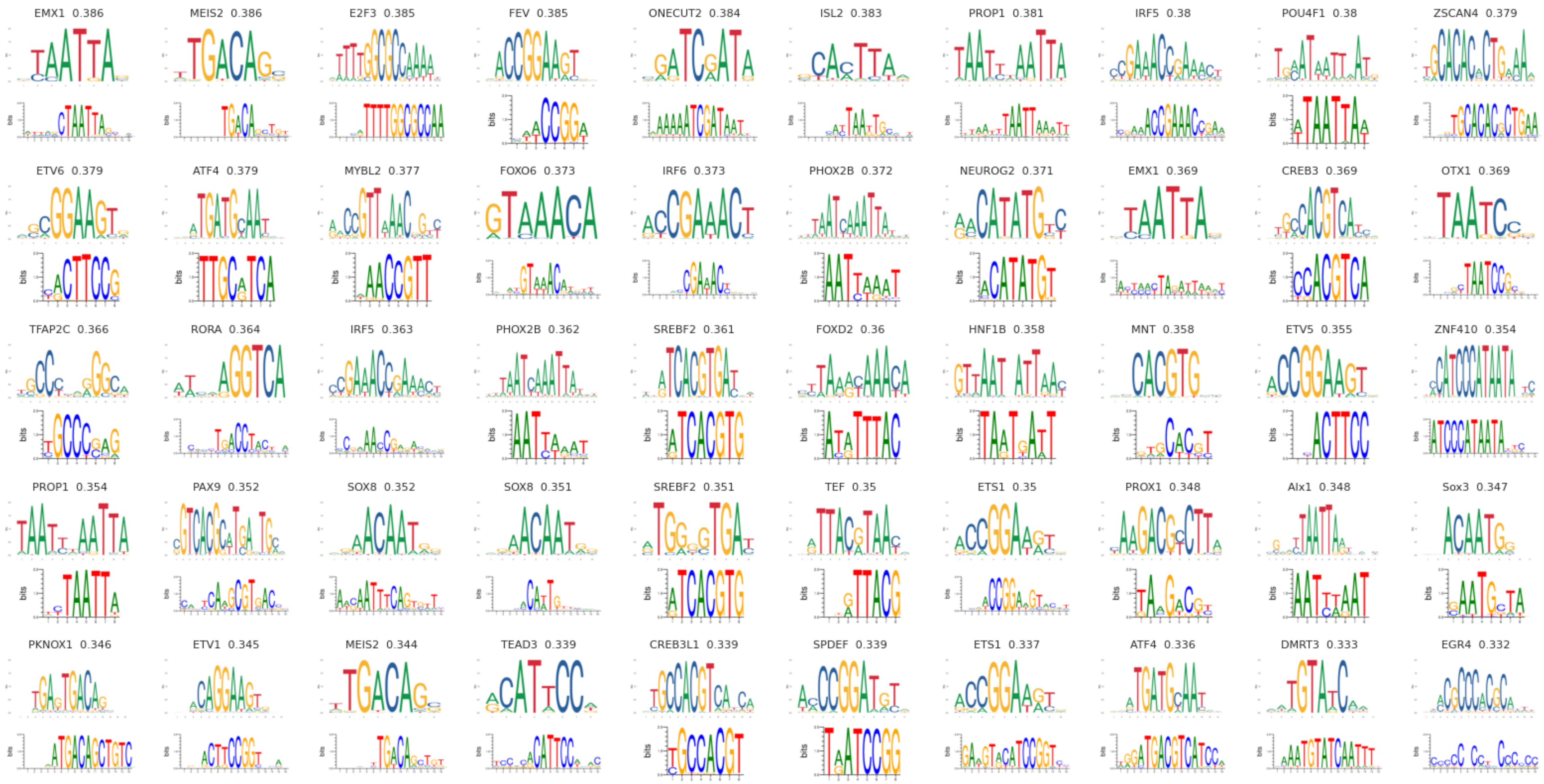


**Figure S9.** The logos of the motif pairs whose similarities ranked from 201 to 250.

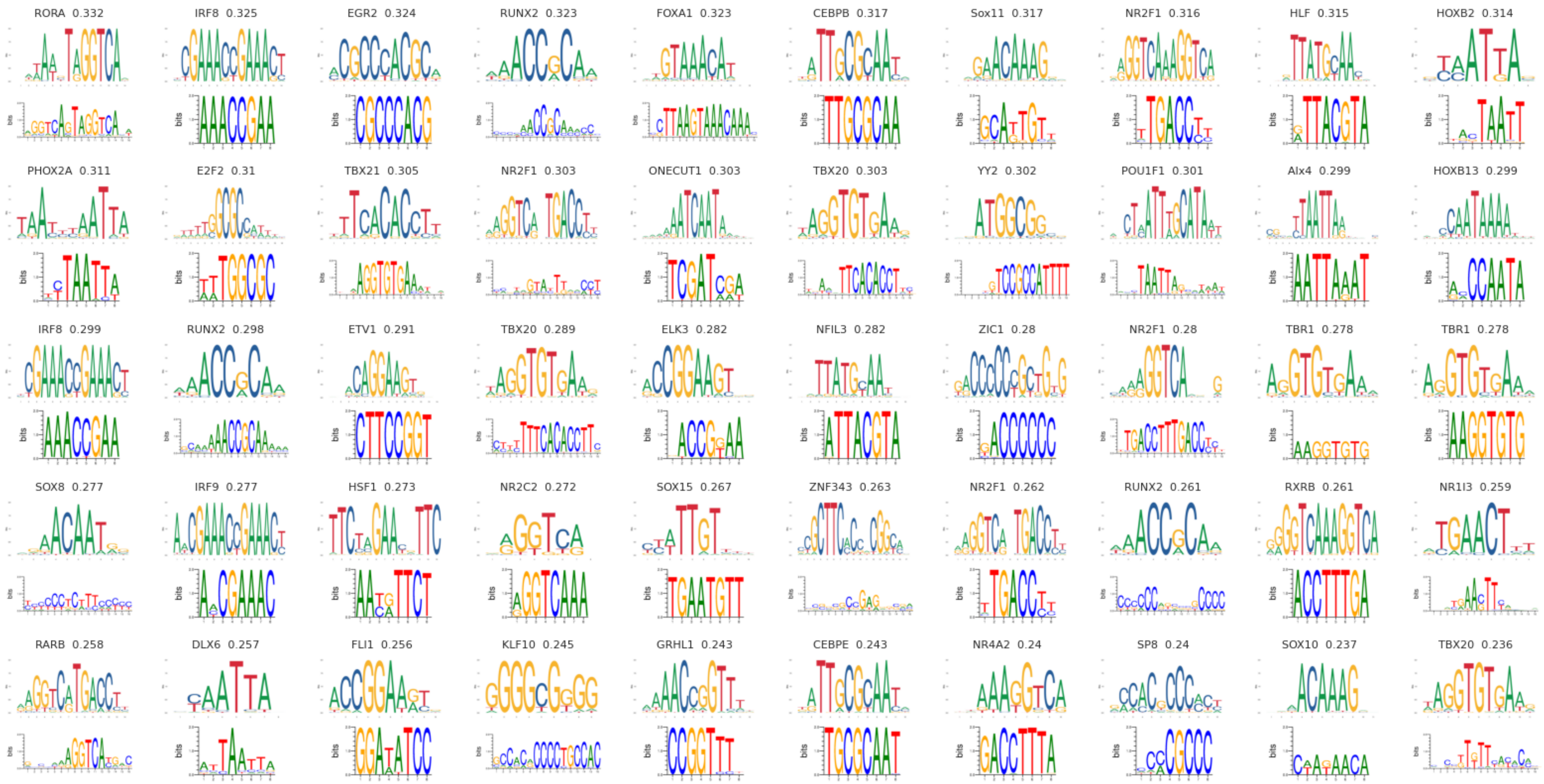


**Figure S10.** The logos of the motif pairs whose similarities ranked from 251 to 300.

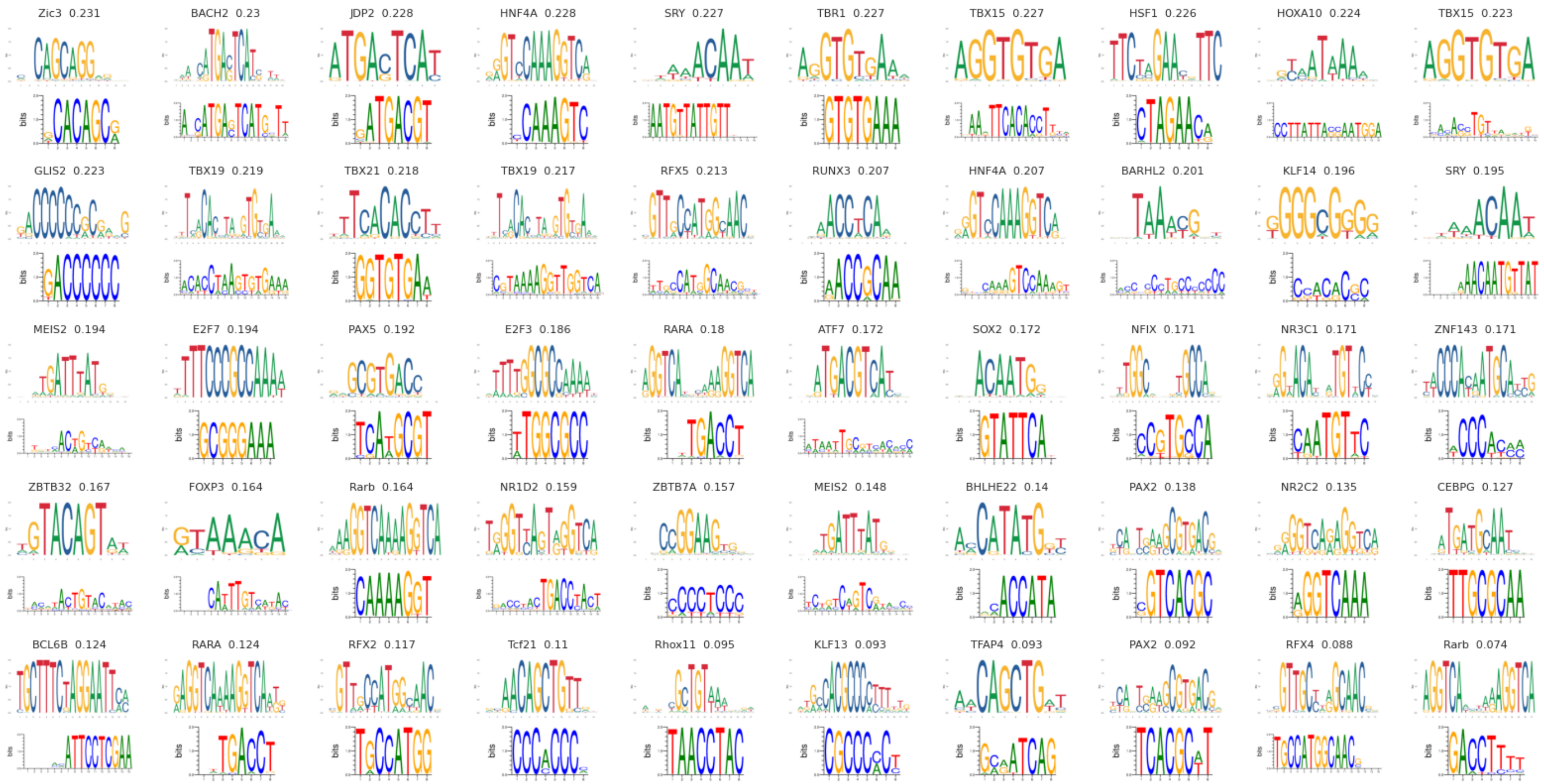


**Figure S11.** The logos of the motif pairs whose similarities ranked from 301 to 350.

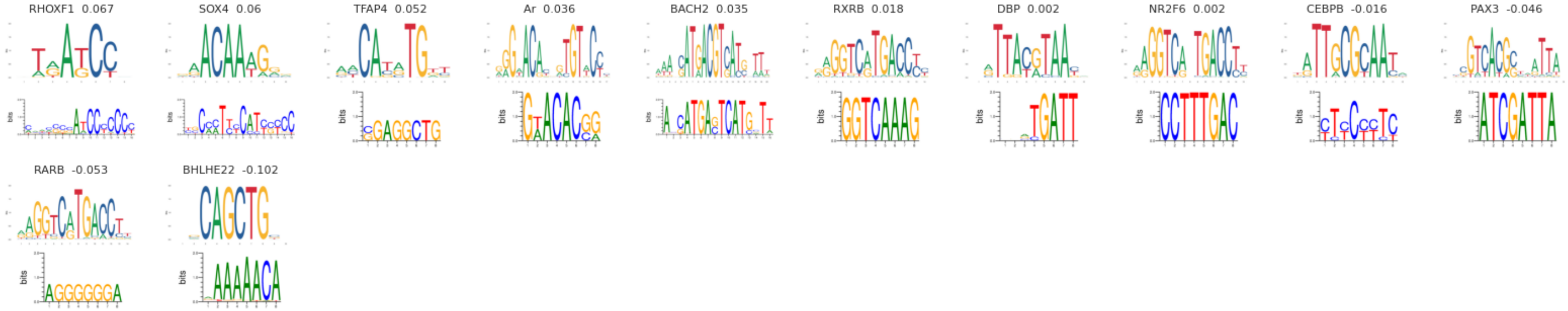


**Figure S12.** The logos of the motif pairs whose similarities ranked from 351 to 362.

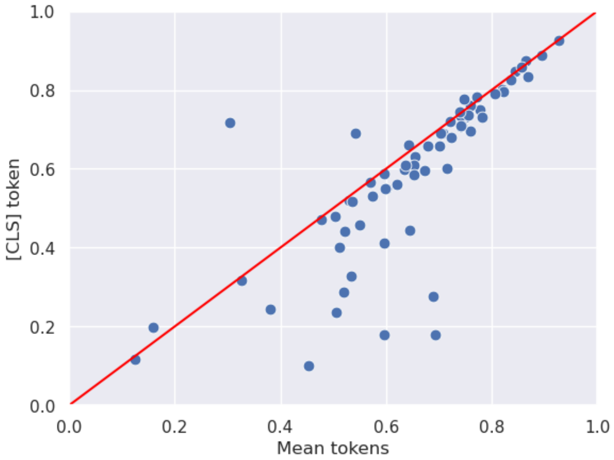


**Figure S13.** The impact of two pooling methods. Each data point was derived from the evaluation results of utilizing two distinct pooling modules.

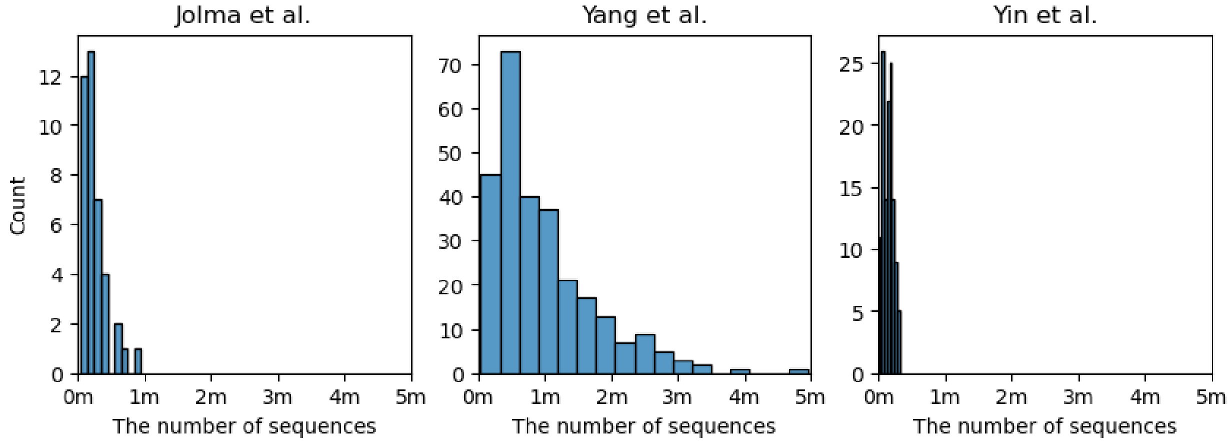


**Figure S14.** The number of sequences on HT-SELEX datasets from three studies. The range of the x-axis is up to 5 million sequences.

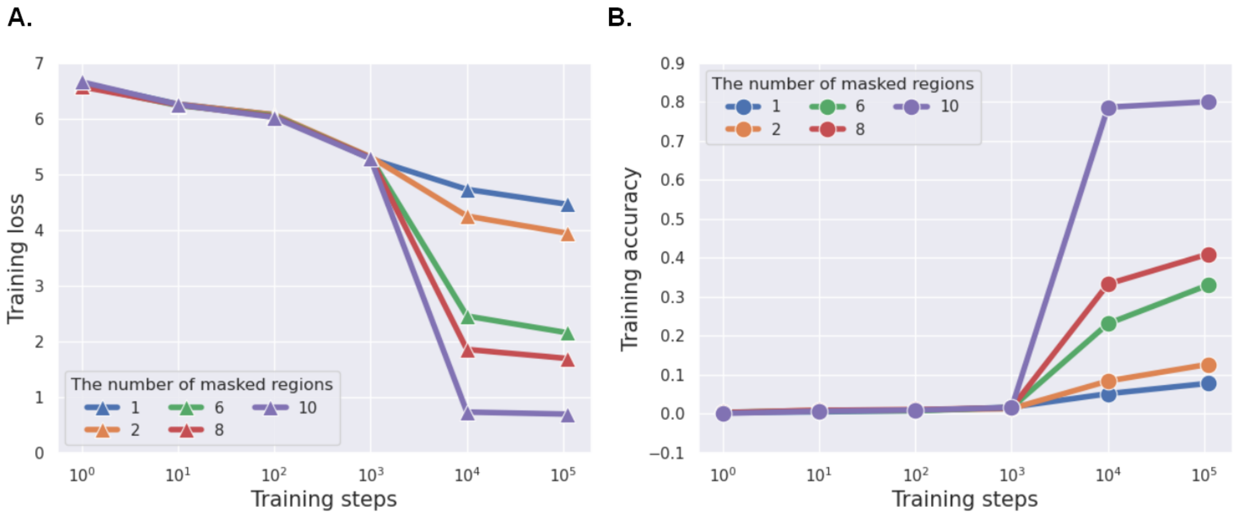


**Figure S15.** The performance of masked token prediction during the pre-training stage. **(A)** The loss of masked token prediction in each pre-training step. **(B)** The accuracy of masked token prediction in each pre-training step.
